# High resolution, proteome-wide mapping of subcellular protein localization in plants

**DOI:** 10.64898/2026.02.27.708449

**Authors:** Monique van Schie, Mark Roosjen, Catherine Albrecht, Jarik van Marsdijk, Dolf Weijers

## Abstract

Protein function is intimately connected to subcellular localization, and experimental determination of protein localization is a key element of understanding biological roles. However, even in the best-studied model plants, such as *Arabidopsis thaliana*, a minority of proteins has an experimentally defined subcellular localization. We present an experimental strategy to globally map plant subcellular proteomes by mass spectrometry. We annotated subcellular localization of 7815 proteins in Arabidopsis roots, 4672 in Arabidopsis seedlings, and 2782 in the liverwort *Marchantia polymorpha*. By independent validation, we find that these annotations are highly predictive and can be integrated with other proteomics datasets. Cross-species comparisons reveal substantial global conservation of subcellular localization. Furthermore, we demonstrate that the same approach can be used to identify dynamically translocating proteins upon treatment or in a mutant. This work shows the power of global spatial proteome mapping in plants and offers an extensive resource for protein subcellular localization in plants.

**Highlights:**

- Optimized approach for global mapping of protein subcellular localization by differential centrifugation in plants
- Interactive resource of subcellular localization of plant proteins at unprecedented depth and resolution
- Cross-species comparison reveals that the plant subcellular proteome is deeply conserved
- Comparative subcellular proteomics of a Brefeldin A treatment and a *gnom* mutant robustly describes global shifts in protein localization

## Introduction

In eukaryotic cells, the subcellular localization of proteins is tightly connected to their function. The different microenvironments within the various compartments of the cell not only offer optimized conditions for specific protein functions, but also offer the potential to concentrate or separate proteins^1^. Hence, translocation of proteins is often involved in biological processes^2^. Studying subcellular localization and protein translocations at a cell-wide scale is therefore crucial to our understanding of protein functions and their role in cellular processes^3^.

Over the past decades, numerous approaches have been developed to study subcellular localization of proteins. Among these are microscopy-based methods that require the fluorescent tagging of proteins, or the use of antibody staining, offering high precision but low throughput^4^. Isolation of single organelles followed by mass spectrometry is also widely used, which offers a higher throughput compared to microscopy, but often proteins from other organelles are co-purified^5^. More recently, proximity ligation has been proven to be a powerful tool to identify proteins in cellular niches^6,7^. These methods all offer specific benefits, but do not provide a global overview of the cell.

To obtain this global overview of protein subcellular localization, multiple methods have been established that rely on differential protein profiling. All these approaches are based on the assumption that proteins will behave very similar if they are derived from the same subcellular location^8–11^. In general, a gentle native lysis is followed by a separation of the cellular compartments based on either physical or chemical properties. Separation is for example achieved based on relative weight by the application of differential centrifugation^8^ or density gradients^9^. Furthermore, separation based on differences in chemical properties^10^ or by the use of labeled proteins pulled down with antibodies^11^ have been demonstrated. These efforts have yielded many new biological insights, for example on protein trafficking related diseases and cellular responses to viral infections^12,13^.

For plants, information on subcellular localization of proteins is more fragmented. There are several examples of proteomes of individual organelles such as mitochondria^14–18^, vacuoles^19–21^, chloroplasts ^22,23^, the endomembrane system^24–28^, cytosol^29^, and nucleus^30,31^, but there is only a single example of a global profiling of subcellular localization that encompasses 527 Arabidopsis proteins^32^. Hence, there is little insight in global plant proteome organization, likely mostly because of challenges posed by the very nature of plant tissues. Plants have rigid cell walls and large vacuoles, which makes controlled cell lysis to obtain intact organelles more challenging^33^. Furthermore, plant proteins are generally not as well annotated compared to those in other model species, hence identification rates tend to be lower and as a consequence, there are fewer well-validated subcellular marker proteins^34^.

We here describe a strategy building on existing protein profiling methods based on differential centrifugation approaches, which we optimized for use on plant tissues. We optimized the complete workflow including sample preparation, data acquisition and downstream data processing and analysis. We present a resource deep subcellular proteome of the flowering plant *Arabidopsis thaliana*, both at the organ level for roots and the organismal level for seedlings. Furthermore, we compare the Arabidopsis subcellular proteome to that of the liverwort *Marchantia polymorpha*, two species that diverged around 430 million years ago, and which allowed us to study the evolution of protein subcellular localization in plants^35,36^. In addition to the analysis of static subcellular proteomes, we also demonstrate that translocating proteins can be identified on proteome-wide scale by performing a Brefeldin A (BFA) treatment and studying the trafficking mutant *gnom*^37^. Lastly, all the presented data, including the translocation data sets, is interactively browsable in a set of Shiny apps through the following links: [Roots](https://weijerslab.shinyapps.io/spatial_proteomics_roots_arabidopsis_thaliana/), [Seedling](https://weijerslab.shinyapps.io/spatial_proteomics_seedling_arabidopsis_thaliana/), [Marchantia](https://weijerslab.shinyapps.io/spatial_proteomics_marchantia_polymorpha/), [BFA treatment](https://weijerslab.shinyapps.io/spatial_proteomics_dynamic_bfa_treatment/), [gn-fwr](https://weijerslab.shinyapps.io/spatial_proteomics_dynamic_gnfwr/).

## Results

### An optimized procedure for global mapping of subcellular protein localization in Arabidopsis

No established methods are yet available for deep, global mapping of subcellular protein localization in plants. We therefore compared two similar methods that are commonly used in other eukaryotic species for their applicability to plants^8,38^. Both Localization of Organelle Proteins by Isotope Tagging after Differential Centrifugation (LOPIT-DC) and Dynamic Organellar Maps (DOMs) methods are based on the separation of organelles by differential centrifugation, followed by protein identification and quantification in the individual fractions by mass spectrometry^8,38^. The underlying principle is that instead of isolating organelles individually, the cell contents are fractionated in a pattern where every organelle or compartment displays a unique distribution pattern over the fractions. These patterns can then be used to cluster co-localizing proteins using statistical methods, to then determine the identity of these clusters through the use of organellar marker proteins with well-established subcellular localizations^8,38^ (Figure 1A).

**Figure 1:**
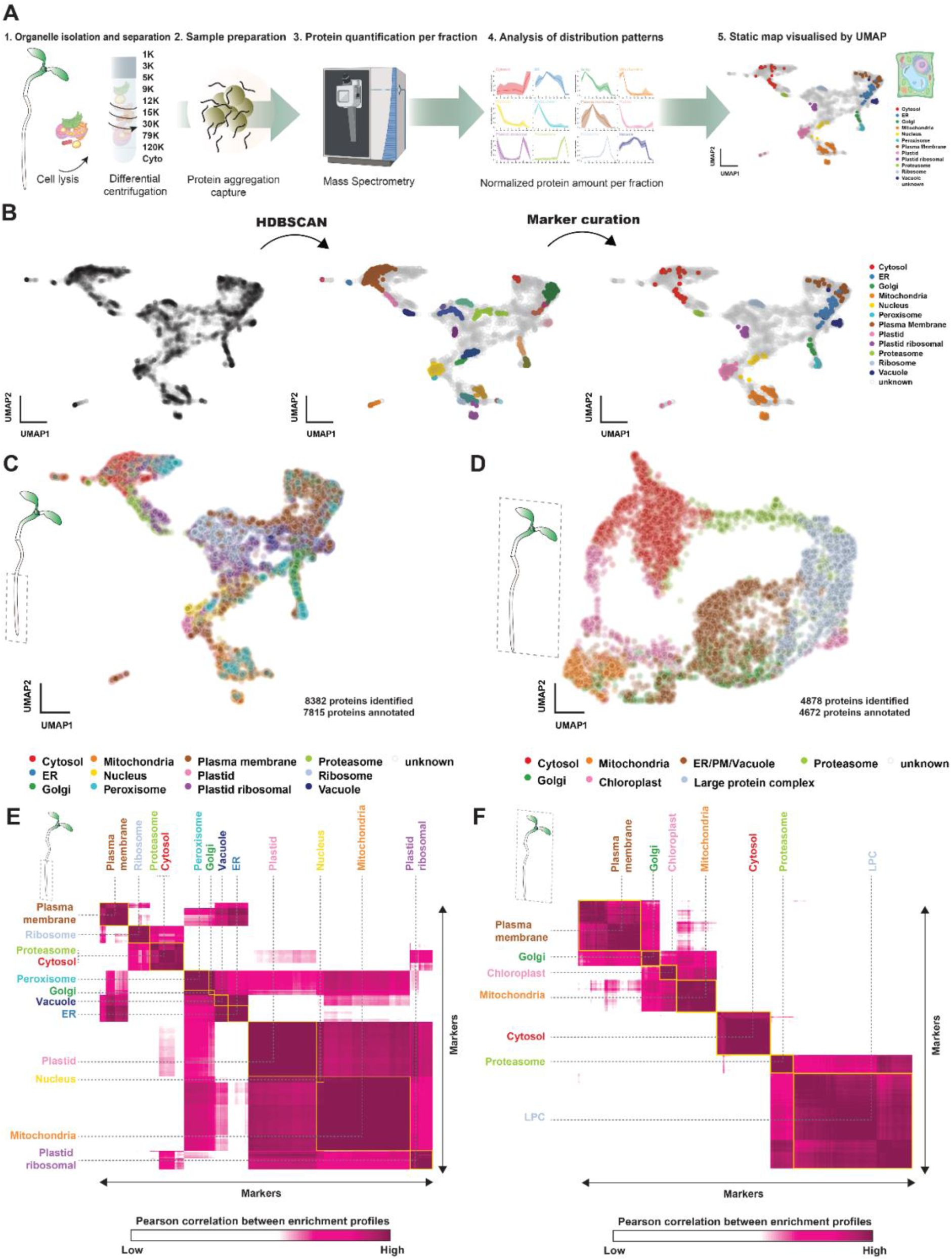
Spatial proteomes of Arabidopsis roots and seedlings. (A) Overview of the method for generating spatial proteomic maps. Plant tissues are lysed and contents are separated by differential centrifugation into 10 fractions. The proteins in each fraction are identified and quantified by mass spectrometry. Protein abundance is normalized over fractions, resulting in unique patterns. All proteins in the dataset are visualized by UMAP based on their fractionation patterns and marker proteins are used to annotate regions of the map. (B) Overview of the procedure to identify the subcellular localization of clusters in the UMAP to curate the marker protein list. First, clusters are identified in a data-driven manner using the HDBSCAN algorithm. Next, proteins in every cluster are checked for experimental evidence of subcellular localization. Lastly, clusters are curated based on the quality of the marker proteins and a list of high-quality marker proteins. (C) UMAP showing the Arabidopsis root dataset with 7815 proteins annotated to 12 subcellular localizations. (D) UMAP of the Arabidopsis seedling dataset with 4672 proteins annotated to 7 subcellular localizations. (E) Pairwise Pearson correlation of the fractionation patterns of marker proteins for the root dataset. There is a strong correlation within the same subcellular localization, but for some localizations there is also a lower correlation with different subcellular localizations. (F) Pairwise Pearson correlation of the fractionation patterns of the marker proteins for the seedling dataset. There is a strong correlation within the same subcellular localization.

The critical parameter in obtaining reliable subcellular spatial proteome maps is complete lysis of tissues and cells, while retaining organelle integrity during the lysis and centrifugation. Particularly the presence of connecting cell walls in multicellular plant tissues is a major impediment in achieving this. We therefore first explored cell lysis conditions and tested grinding of fresh tissue and automated mechanical homogenization of fresh tissue, each followed by potter homogenization. We tested the integrity of organelles by Western Blots with organellar marker antibodies and found that automated mechanical homogenization offers the best balance between retaining organelle integrity and reproducibility (Figure S1).

We next compared the LOPIT-DC (10 fractions) and DOMS (6 fractions) methods in their ability to separate plant cell compartments. We separated root lysates by differential centrifugation according to each method, and quantified proteins in each fraction by nano-Liquid Chromatography coupled to tandem Mass Spectrometry (nLC-MS/MS). Using an objective measure for separation of markers and compartments (mean profile difference and average distance between clusters), we found that LOPIT-DC allowed better reproducibility of the fractionation patterns (Figure S2). To maximize the number of protein identifications and quantifications, we implemented Data-Independent Acquisition (DIA) and compared this analysis mode to Data-Dependent Acquisition (DDA) on the same samples. We found DIA to increase the identified number of protein groups by 60%, while also improving the reproducibility of the data (Figure S2).

Having established a reliable lysis and fractionation method for Arabidopsis roots, we next explored data analysis and visualization. Normalized MS quantifications for each protein across all fractions of three replicate experiments were projected on a UMAP^39^ (Figure 1B). Next, clusters of proteins with similar distribution patterns were statistically identified by the algorithm HDBSCAN^40^ (Figure 1B). Afterwards, the identity of each cluster was determined using well-established marker proteins, based on experimentally validated microscopy localizations in the SUBA5 database^41^, supplemented by experimentally derived information from the Uniprot database^42^ where necessary. Only proteins with experimentally derived localization to a single organelle were selected to be a marker. Clusters that did not show a clear consensus on their subcellular localization were omitted from further analysis and markers from clusters with the same consensus localization were merged to one localization in the final marker list. Finally, we used a Bayesian T-Augmented Gaussian Mixture model Markov-chain Monte-Carlo (TAGM-MCMC)^43^ to predict the non-marker proteins and visualized these in a UMAP.

The resulting Arabidopsis root subcellular proteome map (Figure 1C) contains 8382 proteins, of which 7815 proteins could reliably be classified to a subcellular location (TAGM-MCMC probability score >0.999 and TAGM-MCMC outlier score <5e-5). Next, following the same procedure (with a minor modification of the homogenization, see STAR★Methods), we generated a second spatial subcellular proteome map from entire Arabidopsis seedlings. In this seedling dataset, we measured 4878 proteins and assigned 4672 proteins to a subcellular localization. The correlation of the distribution patterns for the various marker proteins confirm that we have sufficient separation and that these patterns are reproducible (Figure 1E,F). The distribution profiles of the proteins over the fractions also match the expected patterns, where relatively heavy organelles such as the nucleus and ER peak in the early fractions, while lighter compartments such as the cytoplasm and proteasome peak in the latest fractions (Figure S3).

The root map has a higher resolution compared to the seedling map, presumably because the tissue is more homogeneous. Furthermore, the root data set is also deeper, most likely because the chloroplast proteins in the seedlings are so highly abundant that they mask lower-abundant proteins. When we quantify the distribution of proteins over the organelles, we see that the cytoplasm cluster contains most proteins, followed by membrane compartment clusters. The mitochondrial and chloroplast clusters also contain a significant portion of the proteins, while clusters encompassing smaller organelles such as peroxisomes contain less proteins (Figure S3). In summary, we have established a reproducible and sensitive approach for global mapping of subcellular spatial proteomes in Arabidopsis.

### Subcellular spatial proteome maps accurately predict *in vivo* protein localization

Most proteins in Arabidopsis have no experimentally validated subcellular localization, and the subcellular spatial proteome maps can in principle assign high-confidence subcellular localization to thousands of proteins, provided that the inferred patterns reflect true *in vivo* localization. To test the accuracy of the annotations made based on subcellular spatial proteome maps, we validated a selection of annotations by confocal microscopy. From the root spatial proteome map, we selected 35 proteins for which no experimentally defined subcellular localization has been reported (in the SUBA5 database^41^), and chose these such that they would broadly represent the range of subcellular localizations (Figure 2A). Among the selection, most of the proteins have an allocation probability close to 1, and some proteins were included that are predicted to localize to multiple locations (Table S1).

**Figure 2:**
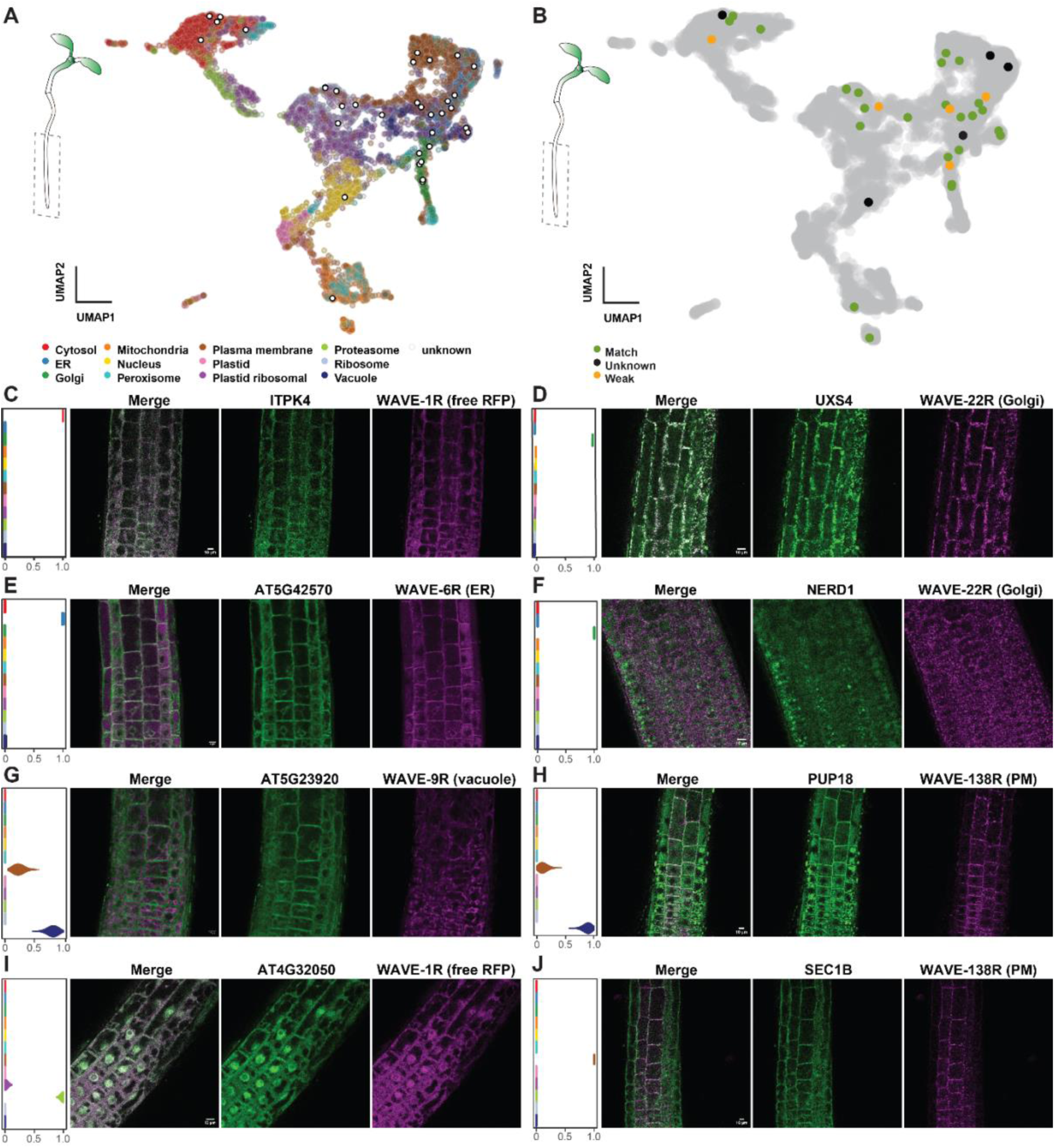
Independent validation of inferred protein localizations in spatial proteomes. (A) The 35 selected proteins annotated with white dots on the root UMAP. (B) Annotation of matches between prediction and microscopy-based localizations. Of the 35 proteins, 25 had a good match (green), 5 a weak match (orange) and 5 proteins had an unknown localization because of low expression levels (black). (C-J) Confocal imaging of selected proteins with a good or weak match to the inferred localization. In each panel, the localization inferred from the map is indicated in the violin plot at the left, with each localization identity color-coded as in (A). (C) ITPK4 in WAVE1-R background [good]. (D) UXS4 in WAVE22-R background [good]. (E) At5G42570 in WAVE6-R background [good]. (F) NERD1 in WAVE22-R background [weak]. (G) At5G23920 in WAVE9-R background [good]. (H) PUP18 in WAVE138-R background [good]. (I) At4G32050 in WAVE1-R background [good]. (J) SEC1B in WAVE138-R background [good]. Scale bar is 10 µm in (C-J).

All selected proteins were C-terminally fused to an mTurquoise2 fluorescent protein, driven by a *UBQ10* promotor and stably transformed into Arabidopsis (Figure 2; Figure S4; Table S1). To determine the subcellular localizations more accurately, where available, we selected established organellar marker backgrounds in which each construct was transformed^44,45^ (Table S1). We next analysed subcellular protein localization in transgenic roots relative to these markers, and scored localization to be either a good match, a poor match, no match, or unknown. Given that we used an overexpression promoter, the match of the observed pattern with the predicted one was considered good if there was co-localization, even if experimental localization was broader. We consider proteins that are found in similar organelles a weak match and scored no match if there is no overlap at all. We visualised the set accordingly and plotted the results on the root map (Figure 2B; Table S1).

Of the 35 imaged proteins, 30 could be detected at levels that allowed for determining localization. Of these 30, there were 25 with a good match to the inferred pattern, while the remaining 5 showed a weak match (Figure 2C-J; Figure S4; Table S1). Importantly, there was no clear systematic bias in the degree of positive validation depending on the predicted localization. This shows that annotations based on the root map can be made with high accuracy (84%, based on 30 proteins) across various subcellular localizations (Figure 2B).

Most proteins included in this validation series had a single predicted location, while some had dual predictions. For example, two proteins are predicted to reside both in the vacuole and the plasma membrane, and we find support for both locations in the imaging data (Figure 2G,H). This indicates that more complex patterns can also be validated and thus that the subcellular spatial proteome maps can accurately annotate the subcellular localization of previously undescribed proteins.

### Integration of subcellular spatial proteome maps with complementary proteomics datasets

Having confirmed that the subcellular spatial proteome maps can successfully and globally annotate subcellular protein localization, we explored the potential of the maps further. We tested whether integration of a variety of complementary datasets with the Arabidopsis root map can generate new biological insights or help generate new hypotheses, by studying protein complexes, liquid-liquid phase separation and protein-protein proximity proteomics as proof of concept.

Proteins must colocalize to interact, so obtaining large scale data on co-localization would be a valuable resource to identify new potential protein interactions and complexes. However, it is unclear if the methodology used to generate spatial subcellular proteome maps leaves existing protein complexes intact. We explored this question by mapping a previously reported census of stable protein complexes^46^ (Figure 3A) onto the root map. We first asked whether the known members of protein complexes are all annotated in the same subcellular location. As expected, many complexes in the root can be found in the cytosol. Mitochondrial and chloroplastic ribosomes can be found in their respective organelles. While we see that most members of known complexes cluster in the map, for some complexes such as the AP1/2 complex, not all members cluster in the same region of the map. We conclude that in general, protein complexes stay associated during the procedure. We next asked if the distance between proteins in the UMAP could be leveraged to identify new protein complexes. We compared the average Euclidean distance between proteins from each complex to an equivalent random draw of proteins from the respective organelle. In this comparison, we found no significant difference between the median pairwise correlation between protein complexes and a random draw from an organelle (Figure 3B). Therefore, while the maps reflect the co-localization of known protein complexes, they cannot be used to infer new protein interactions, likely because the fractionation profiles are based on the separation of complete organelles.

**Figure 3:**
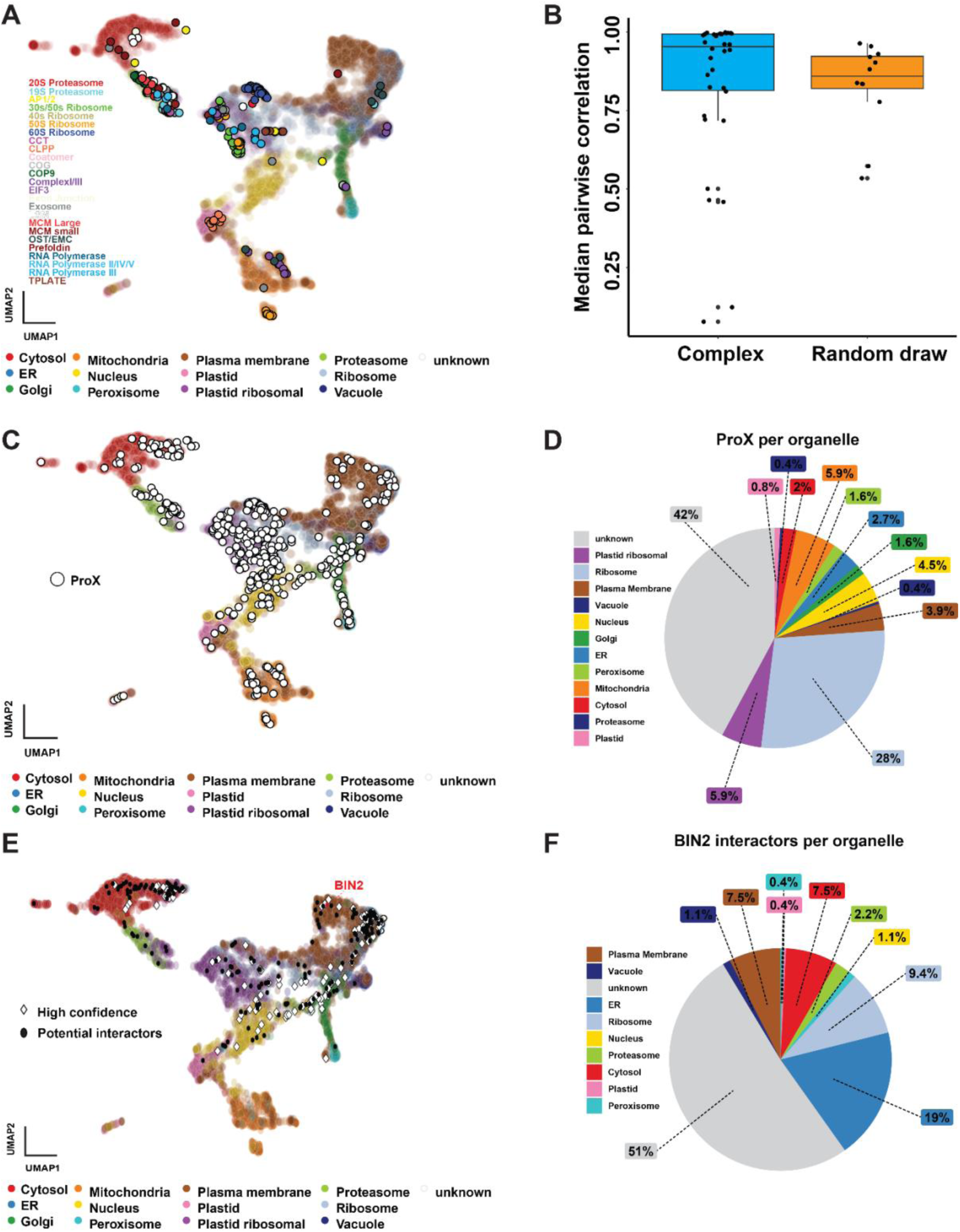
Integration of spatial proteomes with other datasets. (A) Protein complexes identified by McWhite et al.^46^ projected on the root UMAP. (B) Comparison of the average Euclidean distance between proteins from each complex to an equivalent random draw of proteins from the respective organelle showing no significant difference, indicating that no new protein complexes can be inferred using the root subcellular proteome data. (C) A census of Arabidopsis proteins prone to phase-separate based on precipitation by b-isox crystals (ProX)^47^ plotted on the root map. (D) Distribution of (C) as percentage of the subcellular localizations shows high enrichment in the ribosome cluster and unknown regions of the map. The quantification also shows low enrichment in the membranous compartments, suggesting that protein condensates may display unique patterns in the root dataset. (E) Proximity labelling dataset using the kinase BIN2 as a bait^48^ projected on the root UMAP, high confidence interactors (diamonds) were identified based on phosphoproteomics, downregulation during bikinin treatment and responsiveness to BR. Two clusters of high-confidence interactors can be identified in the cytosol and the plasma membrane, respectively. (F) Distribution of subcellular localizations of potential BIN2 interactors.

Cells are organized through internal compartments (organelles), large protein complexes, and through the liquid-liquid phase separation of proteins in membraneless organelles. We asked whether such membraneless organelles can be identified in the maps. A census of Arabidopsis proteins prone to phase-separate has been reported based on precipitation by b-isox crystals (ProX)^47^ and we plotted this set of proteins on the root map (Figure 3C). We find these proteins to be highly enriched in the “ribosome” cluster, as well as in unknown regions of the map, while they are almost absent in the membranous compartments (Figure 3D). This suggests that protein condensates may have unique sedimentation behavior that separates them from traditional protein complexes and organelles, yet that resembles the behavior of ribosomes. The parts of the map that is enriched for these proteins may thus offer a good starting point for identifying novel phase-separating proteins.

Many studies use protein-protein interaction screens to understand the context in which a protein acts. In recent years, proximity labeling (PL) has emerged as a powerful approach to map proteins in the vicinity of a protein of interest. However, PL datasets tend to contain substantial false positive hits because PL labeling requires only transient vicinity of proteins. We asked if mapping the subcellular localization of PL partners in relation to the bait protein can offer context to complex interactomes. We took a recently published dataset of the kinase BIN2 as an example, where interactors were not only identified, but also classified as high-confidence based on data integration with phosphorylation state, downregulation during bikinin treatment and responsiveness to BR^48^. In the root map, BIN2 is localized in the plasma membrane, as are 7.5% of its interactors (Figure 3E, F). We also found interactors in the ER (19%) and cytoplasm (7.5%), as well as a large unknown fraction of 51%. BIN2 has been experimentally localized to the plasma membrane, cytoplasm and nucleus, so this could explain the wide distribution of subcellular localizations of the interactors^49,50^. Furthermore, there are two clusters of high-confidence interactors, one in the cytoplasm and one in the plasma membrane, which could be interaction hubs. The spatial mapping of this PL interactome dataset thus helps identify candidate interactors mediating kinase function in specific compartments (e.g. plasma membrane).

In summary, integration of complementary proteomics data with the subcellular proteome maps offers an extra layer of information that can be leveraged to obtain biological insights.

### Subcellular proteome organization is conserved among land plants

Protein function is deeply intertwined with subcellular localization. During evolution, and through the expansion of gene families, protein localization patterns may change, creating species-specific proteome organization. To explore the extent of conservation and diversification in subcellular proteome organization among land plants, we generated a spatial proteome map of the non-vascular liverwort (bryophyte) species *Marchantia polymorpha*. The last shared common ancestor of Arabidopsis and Marchantia lived around 430 million years ago^51^ and these two species have vastly different life histories and morphologies. Furthermore, Marchantia has recently become a popular model species for bryophyte biology and for comparative plant biology, and hence insight in its protein subcellular localization will be a valuable resource.

To establish the Marchantia thallus subcellular spatial proteome, we used the same protocol and data-analytical pipeline as we used for the Arabidopsis seedling samples (Figure 1A; STAR★Methods). The resulting subcellular proteome contains 3630 proteins, and 2782 proteins were confidently annotated to a subcellular localization (Figure 4A). For the validation of the map, we selected a number of proteins for *in vivo* validation of the localization annotations by confocal microscopy (Figure 4B-H; Figure S6; Table S2). We found that 9 of 14 (64%) of the predictions were validated by localization of fluorescently tagged proteins, which was less accurate compared to the 84% accuracy in the Arabidopsis roots (Figure 2B; Table S1). However, the sample size of imaged proteins is also smaller and spans a smaller range of subcellular localizations.

**Figure 4:**
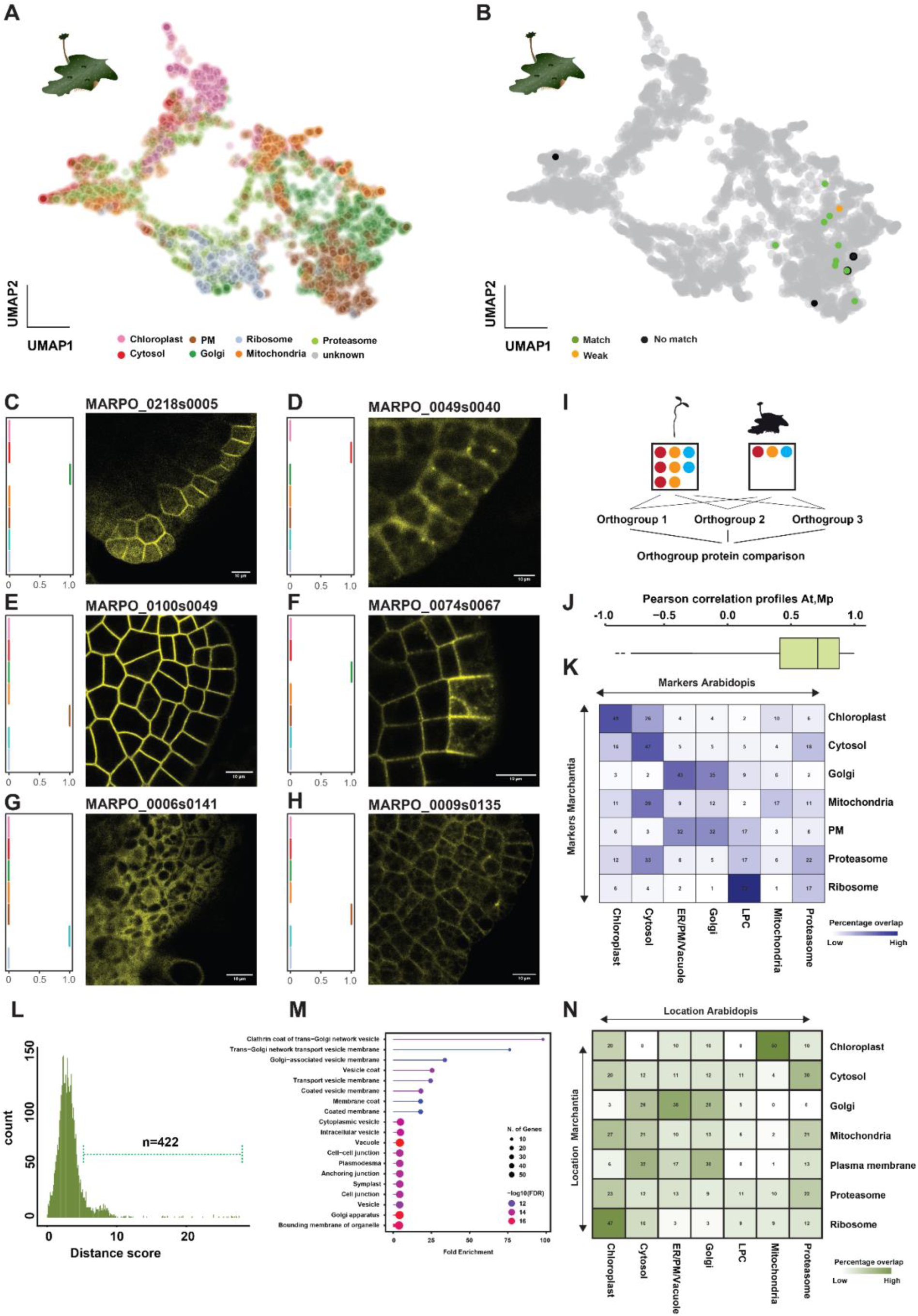
A *Marchantia polymorpha* spatial proteome reveals high conservation of subcellular protein localization. (A) Two-dimensional UMAP of the Marchantia thallus subcellular proteome, with 2782 proteins annotated to 7 subcellular localizations. (B) Marchantia UMAP with selection of 14 proteins for microscopy localization. Of the 14 proteins, 9 had a good match (green), 1 a weak match (orange) and 4 proteins had no match (black). (C-H) Confocal imaging of MARPO_0218s0005 [no match] (C), MARPO_0049s0040 [good] (D), MARPO_0100s0049 [good] (E), MARPO_0074s0067 [weak] (F), MARPO_0006s0141 [no match] (G), and MARPO_0009s0135 [good] (H) in Marchantia gemmae. The inferred localization is shown in the violin plots in the left of each panel and color-coded for clusters as in (A). (I) Construction of orthogroups between *Arabidopsis thaliana* and *Marchantia polymorpha*. (J) Pearson correlation of the fractionation patters of the proteins in the same orthogroup, showing a high conservation of 0.59 on average. (K) Overlap of protein markers for subcellular localization between Arabidopsis and Marchantia, confirming a high conservation of protein subcellular localization. (L) Distribution of evolutionary distance score of proteins from the same orthogroup, based on distance in a joint UMAP. Most proteins have a low evolutionary distance score, so their subcellular localization is likely conserved, while 422 proteins have a high evolutionary distance score. (M) GO-term analysis of the 422 proteins in (L) with a high Evolutionary Distance score. (N) Comparison of the subcellular localization of the 422 proteins with a higher Evolutionary distance score in the Marchantia thallus and Arabidopsis seedling maps. Scale bar is 10 µm in (C-H).

We next compared global proteome organization in the Marchantia thallus map with that of Arabidopsis seedlings. Marchantia does not have tissue that is homologous to roots, whereas thallus and cotyledon are both photosynthetic tissues. We used orthogroups that were previously obtained using OrthoFinder^52,53^ (Figure 4I), analyzed the fractionation patterns of proteins in the same orthogroup, and calculated Pearson correlation coefficients between species. We found an average Pearson correlation of 0.59 (Figure 4J), signifying an extremely high correlation in overall subcellular proteome organization, which is also reflected in a significant overlap of the marker proteins (Figure 4K). Next, we compared the Marchantia and Arabidopsis seedling data using Aligned-UMAP to assess the distance between proteins from the same orthogroup between the two species. As expected, most proteins from the same orthogroup had small distances between them, hence a low Evolutionary Distance Score (Figure 4L). However, this analysis also identified 422 proteins with a higher Evolutionary Distance Score, which we consider candidates for neo-functionalization based on protein localization. As a first exploration of these proteins with a higher Evolutionary Distance score, we performed a GO-term analysis (Figure 4M) and compared the localization of these proteins in Arabidopsis seedlings and Marchantia (Figure 4N). Most of the enriched GO-terms are related to protein trafficking, suggesting that alterations in this process played an important role during plant evolution. We conclude that there is a high degree of conservation on subcellular proteome organization between Arabidopsis and Marchantia, and likely more broadly among land plants.

### Mapping dynamic global spatial proteome remodeling in Arabidopsis roots

The cellular spatial proteome is not static; many proteins dynamically translocate as part of their activity cycle, and protein function is strongly regulated by the dynamic control of localization. We explored whether subcellular spatial proteome maps can report the dynamic localization of proteins on a global scale. We focused on anterograde membrane trafficking, where vesicle transport from ER to Golgi to the plasma membrane controls membrane localization. One of these secretory steps can be inhibited by the well-studied fungal toxin Brefeldin A (BFA)^54–57^. BFA inhibits the function of large ARF-GEF proteins, which are regulators of the small ARF GTPases that recruit the necessary factors for vesicle formation^54,55,58^. The SEC7 domain of large ARF-GEFs interacts with ARF-GDPs for the conversion of the GDP to a GTP, thereby activating vesicle formation. BFA blocks this transition of ARF-GDP to ARF-GTP, causing the ARF-GEF/ARF-GDP complex to get trapped on the membrane. As a consequence, cycling between the membrane and cytoplasm of BFA-sensitive ARF-GEFs and ARF-GDPs is disrupted^58–60^. The vesicle coat proteins, such as clathrin and COPI, are not recruited to the membrane and accumulate in the cytosol^57,61^. Because vesicle formation is disrupted, the TGN and endosomes fuse into aggregates that are commonly referred to as BFA compartments^56^. Interestingly, in Arabidopsis, Golgi proteins do not relocalize to the ER, unlike in other eukaryotes^60,62^.

While many proteins have been shown to change subcellular localization following a BFA treatment in Arabidopsis roots^60,61,63–66^, there is no proteome-wide view on BFA-sensitive trafficking. We therefore prepared Arabidopsis root maps after 30 minutes of treatment with 10 µM BFA, and prepared control maps of roots treated without BFA. Treated and mock-treated tissue was harvested pairwise three times on separate days and lysis, centrifugation and protein measurements were performed according to the established root protocol. For further analysis, the same data pipeline was used to quantify and normalize the amount of protein per fraction and Aligned-UMAPs of BFA-treated and mock roots were generated. We then used the BANDLE R package^67^ to identify translocating proteins, based on Bayesian Inference statistics.

When comparing the BFA-treated maps to the mock maps, we observe a dramatic rearrangement of organellar clusters, where some are even completely shifted. In an Aligned-UMAP, trajectories of translocating proteins can be visualized, for example between the plasma membrane and the cytosol, from the Golgi and peroxisomes to various localizations and from the ribosomes and nucleus to various localizations (Figure 5A-C). We identified 839 proteins that translocated with a differential localization score of 1 using the BANDLE R package^67^ in the total dataset of 6793 proteins that were measured in both maps, which is more than 10% of the total measured number of proteins (Figure 5C,F). We also measured entire shotgun proteomes on the same samples and found that only 7 proteins significantly changed in abundance (Figure 5D), indicating that identified translocations are not caused by protein abundance changes.

**Figure 5:**
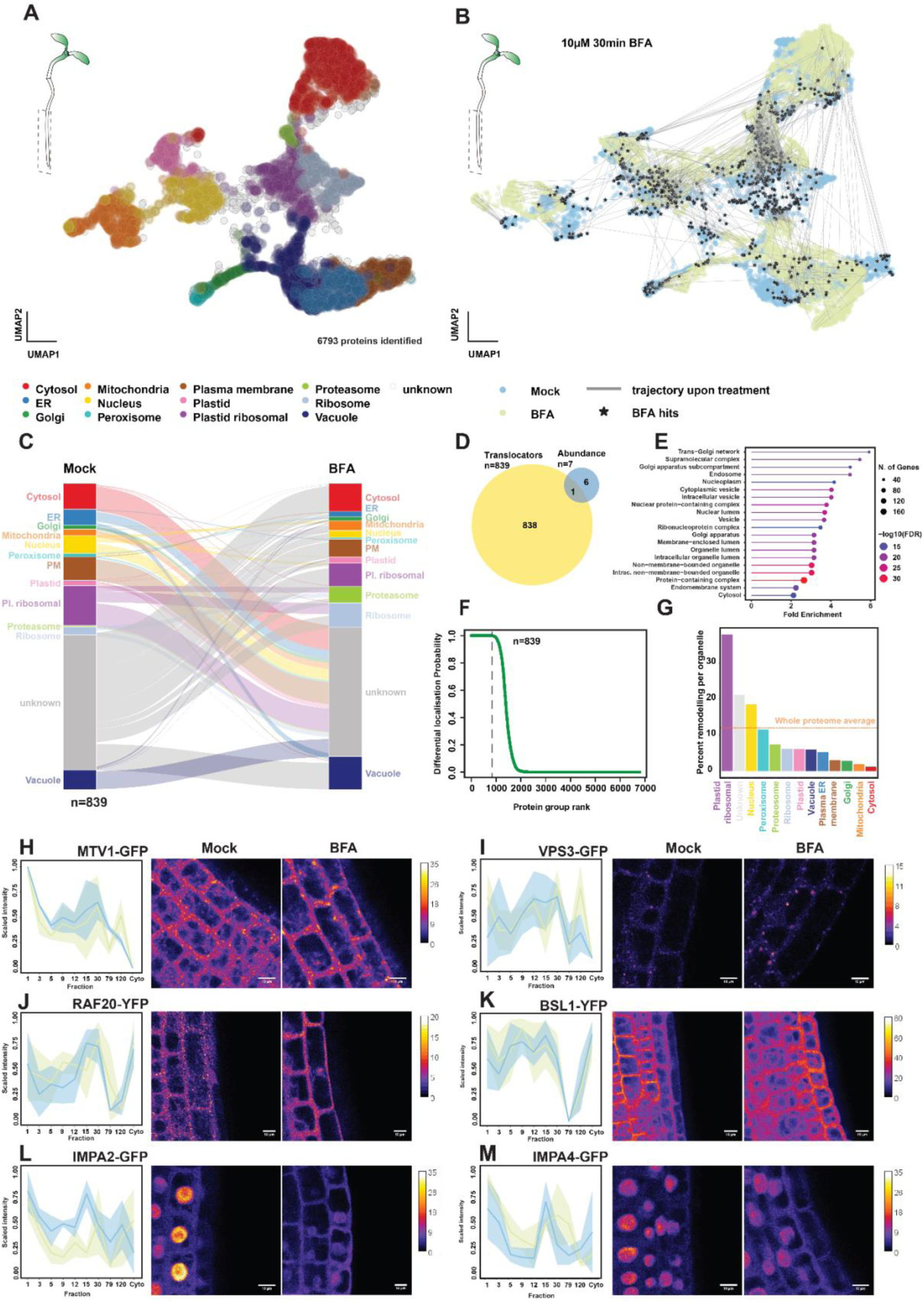
Global subcellular proteome remodeling upon Brefeldin A treatment. (A) UMAP with the subcellular proteome of mock-treated Arabidopsis roots. (B) Aligned-UMAP with the combined data of BFA- and mock-treated roots. Translocating proteins with a differential localisation score of 1 are indicated with stars and their respective trajectories with lines. (C) Alluvial diagram indicating the subcellular localization of the 839 identified translocating proteins in mock (left) and BFA treated roots (right). (D) Overlap between proteins changing in abundance (blue) and translocating (yellow), showing a very small overlap of 1 protein. (E) GO term enrichment for the biological process of the 839 translocating proteins. (F) Distribution of differential localisation probability score of translocating proteins for the whole dataset, in which 839 translocating proteins were identified in a total of 6793 proteins in the dataset. (G) Relative remodeling of subcellular localizations during the BFA treatment. (H-M) Fractionation patterns and confocal imaging for MTV1-GFP (H), VPS3-GFP (I), RAF20-YFP (J), BSL1-GFP (K), IMPA2-GFP (L) IMPA4-GFP (M) in mock- and BFA- treated conditions. Scale bar is 10 µm in (H-M).

Among the significantly translocating proteins are multiple known BFA-sensitive proteins, such as the ARF-GEFs Brefeldin-A Inhibited Guanine nucleotide-exchange protein 1 (BIG1), BIG 2 and BIG5, and the clathrin-interacting protein EPSIN1^63,65,66,68^. To obtain a global overview of the translocating proteins, we performed a GO-term enrichment analysis (Figure 5E). As expected, many proteins that reside in the Golgi, Trans-Golgi, vesicles, and endosomes are affected by BFA treatment. Global analysis of compartment remodeling shows that the “plastid ribosomes”, “unknown”, “nucleus” and “peroxisomes” clusters were most strongly affected (Figure 5G).

We next asked if the comparison of mock-treated and BFA-treated maps accurately reports the effect on individual proteins. We made a selection of 7 proteins that translocate in diverse ways, but that had not previously been shown to be BFA-sensitive, or not at low BFA concentrations. We tested the effect of a 1-hour BFA treatment on the localization of fluorescently tagged proteins in roots (Figure 5H-M Figure S7; Table S3). We observed the commonly described formation of larger aggregates, which are presumably BFA compartments, in the case of MTV1^65,69^ and VPS3 (Figure 5H,I), which was previously only observed for MTV1, and only at 5-fold higher BFA concentrations^69^. For RAF20^53^, we found that distinct cytosolic puncta resolved upon BFA treatment (Figure 5J). For IMPA2 and IMPA4^70^ the intensity of signal decreased in the nucleus and increased for the cytosol (Figure 5L,M). We could not find clear effects for BSL1 and IYO^71,72^, however for these proteins, the difference in fractionation pattern between mock and BFA treatment is much smaller than for some of the other tested proteins (Figure 5K; Figure S7). This may reflect a limit in what can be distinguished by microscopy. In summary, we find that subcellular spatial proteomics can globally identify translocating proteins following drug treatment.

### A mutant spatial subcellular proteome map identifies genetic effects on proteome organization

A major BFA target in Arabidopsis is the ARF-GEF GNOM, best known for its role in the polar trafficking of the PIN auxin transporters^60,64^. GNOM is localized to the Golgi and plasma membrane, and regulates vesicle trafficking, thus mediating developmental patterning^37,60,73,74^. Given that we identified BFA-dependent global protein translocation, we asked if the comparison of *gnom* mutant and wild-type maps allows to triangulate GNOM- and BFA-dependent target proteins.

In this experiment, we also used a more sensitive mass-spectrometer. All other data in this manuscript was obtained using a Thermo Fisher Exploris 480 mass spectrometer, whereas the *gnom* dataset was measured using a Bruker TimsTOF HT mass spectrometer. The number of quantified proteins in wild-type seedlings increased by 72% in half the measurement time compared to the Arabidopsis seedling dataset (Figure 1D). This is important, since it allows for the measurement of more extensive subcellular mapping experiments, while also identifying more regulatory proteins that tend to be lowly expressed.

Since *gnom* null alleles have extremely strong seedling defects, lacking root and hypocotyl^64,75^, we chose the *fewer roots* (*gn^fwr^*) allele, which carries a point mutation causing reduced GNOM activity^37^. This mutant displays a mild phenotype with fewer lateral roots and fragmentation of the leaf vein network^37,73,76^. We generated maps of wild-type and *gn^fwr^* mutant seedlings. We detected 8041 proteins, and by generating an Aligned-UMAP and using BANDLE software, we confidently identified 1553 translocating proteins (differential localisation score > 0.999) (Figure 6A,B). Within the group of translocating proteins, there was clear enrichment of proteins originating from the Golgi and PM/ER/vacuole clusters (Figure 6C,F), which was supported by GO-term analysis of the translocating proteins that showed enrichment of various membranous compartments (Figure 6E). We again measured whole proteomes and found negligible numbers of proteins that changed abundance (Figure 6D); hence the observed translocations are caused by true changes in subcellular localization.

**Figure 6:**
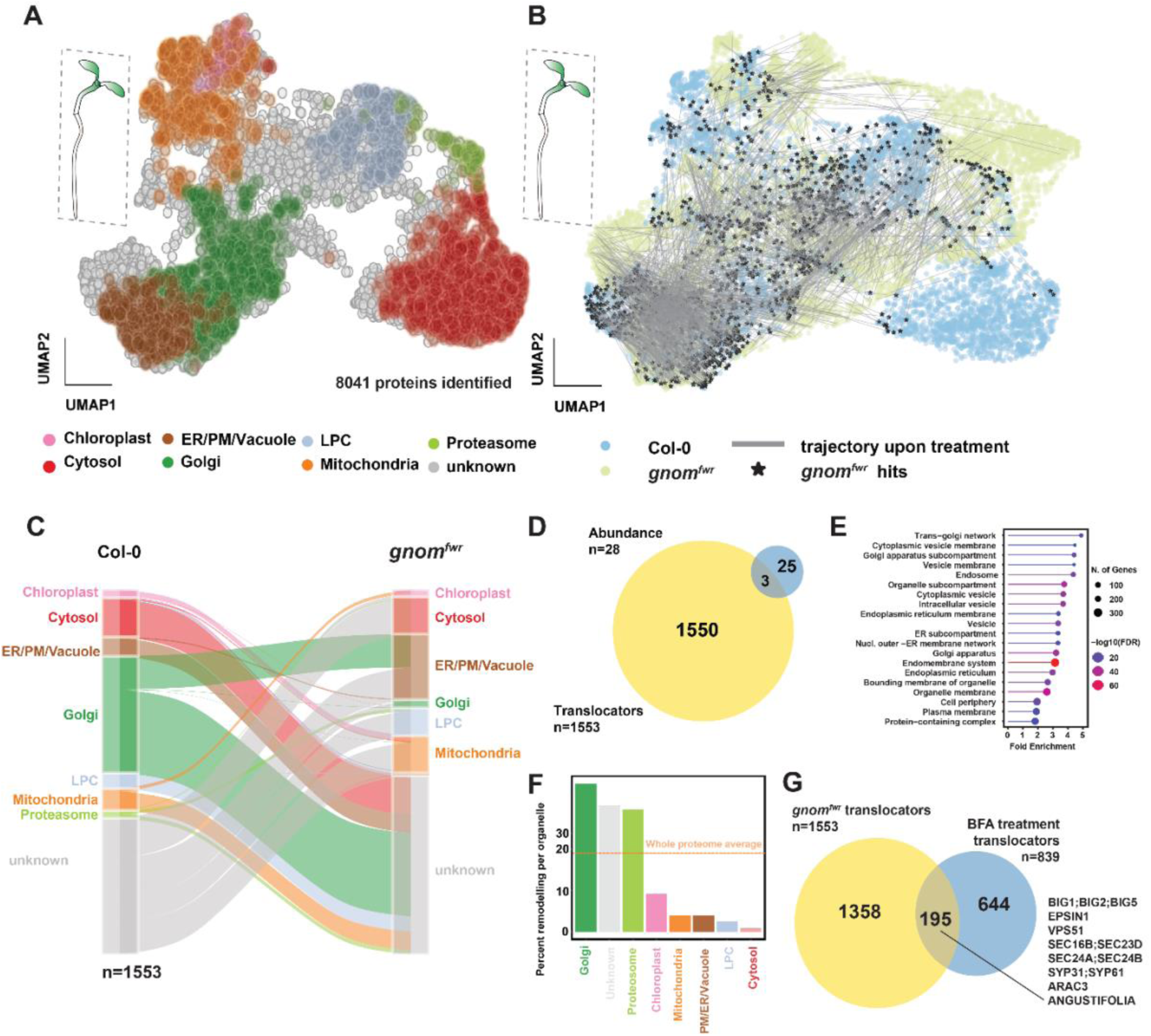
Genetic analysis of subcellular proteome organization. (A) UMAP of Col-0 seedlings. (B) Aligned-UMAP with the combined data of Col-0 wildtype and *gn^fwr^* mutant seedlings. Translocating proteins with a differential localisation probability score of 1 are indicated with stars and their respective trajectories with lines. (C) Alluvial diagram indicating the subcellular localization of the 1553 identified translocating proteins in Col-0 (left) and the *gn^fwr^* mutant (right). (D) Comparison of the number of proteins with significant abundance changes (blue) to the number of significantly translocating proteins (yellow) in the dataset that contains 8041 proteins in total. (E) GO-term analysis of the 1553 translocating proteins. (F) Relative remodeling of subcellular localizations in the *gn^fwr^* mutant compared to the Col-0 wildtype. (G) Quantification of the overlapping significantly translocating proteins between the BFA treatment in Arabidopsis roots (blue) and the *gn^fwr^* mutant seedlings (yellow), showing a significant overlap.

Importantly, detailed analysis of the translocating proteins shows that, as expected, GNOM (as well as its paralog GNL1) is translocating with a score of 1. We identified a number of proteins that have previously been shown to either interact with GNOM or depend on GNOM activity for their localization. These include the GNOM-interacting CYP19-4 (score= 0.916)^77^, Dynamins DRP1a/c and their interactor VAN3 (score=1)^78,79^, PRAF1/4 and RABA2D (score=1)^80^, PIN1 (score=1)^64^ and PIN2 (score=0,988)^81^ auxin transporters, and D6 protein kinase (score=1)^82,83^. Lastly, we found a large number of proteins related to cell wall composition, notably pectin, consistent with cell wall and cell adhesion defects reported in *gnom* mutants^75,84,85^.

Having inhibited the same process both pharmacologically (BFA treatment) and genetically (*gnom* mutant), we asked if the two approaches identify overlapping translocating proteins. Of note, for the *gn^fwr^* mutant, we used whole seedlings, while for the BFA treatment, roots were harvested. Furthermore, the datasets were measured using different mass spectrometers. Finally, the *gn^fwr^*mutant has a developmental phenotype that likely induces indirect protein localization defects, and BFA has other targets beyond GNOM. However, there was still a significant overlap between the translocators of 195 proteins (Figure 6G), which represents 23% of the BFA-induced translocators and 13% of the translocators in the *fwr* mutant. Among these shared translocators, we find BIG1,2,5 and EPSIN, but also more trafficking-related proteins such as vacuolar sorting proteins (e.g. VPS51), SECs and SYPs. We also identified ARAC3, which is involved in PIN2 internalization^86^, and ANGUSTIFOLIA, which has a reported role in the regulation of vesicle trafficking at the TGN^87^.

In summary, mutant spatial subcellular proteome maps report on global changes in protein localization and combining genetic and pharmacological perturbations helps converge on high-confidence target proteins.

### An interactive interface for spatial subcellular proteome maps

This works presents a vast resource of protein subcellular localizations and translocations, which can be mined for multiple purposes. Hence, we decided to integrate all data into interactive Shiny apps^43,67,88^. The datasets described in this study can be found as independent apps: [Roots](https://weijerslab.shinyapps.io/spatial_proteomics_roots_arabidopsis_thaliana/), [Seedling](https://weijerslab.shinyapps.io/spatial_proteomics_seedling_arabidopsis_thaliana/), [Marchantia](https://weijerslab.shinyapps.io/spatial_proteomics_marchantia_polymorpha/), [BFA treatment](https://weijerslab.shinyapps.io/spatial_proteomics_dynamic_bfa_treatment/), [gn-fwr](https://weijerslab.shinyapps.io/spatial_proteomics_dynamic_gnfwr/). In each app, proteins can be browsed based on Uniprot identifiers, Araport codes and gene names. In addition to the placement on the UMAP, it is also possible to study the underlying fractionation patterns. Furthermore, the complete datasets can be downloaded as .txt file and plots of the UMAPs and fractionation patterns as editable .pdf files. In Figure 7, we give several examples of how the data in these apps could be leveraged.

**Figure 7.**
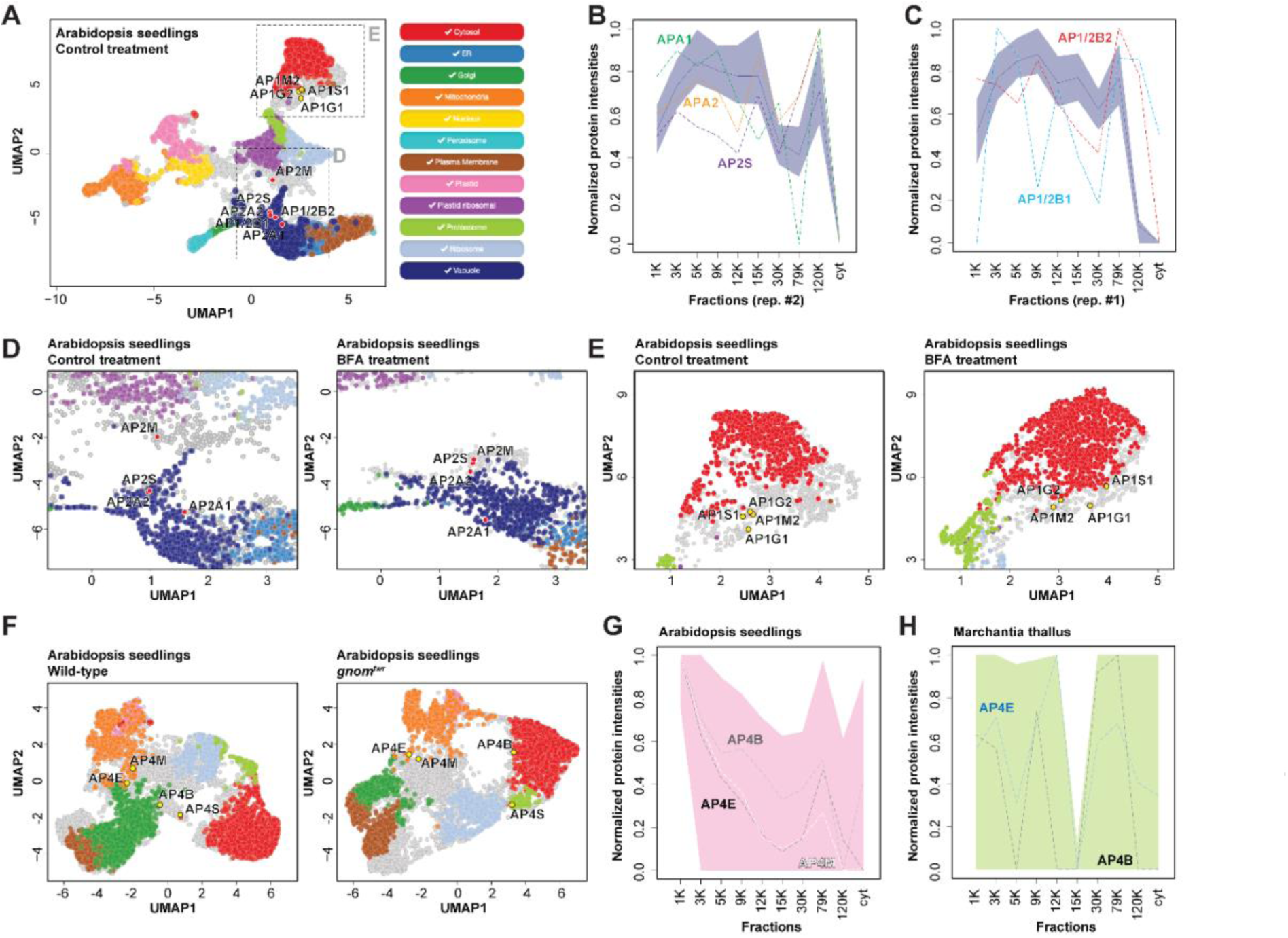
Interactive data exploration with spatial proteomics Shiny apps. (A) Interactive UMAP of mock-treated Arabidopsis roots from the BFA dataset with the AP1 and AP2 complexes annotated. (B) Fractionation patterns of mock-treated Arabidopsis roots from the BFA dataset with the vacuole marker profile indicated in dark blue and the fractionation patterns of the AP2 subunits annotated. (C) Fractionation patterns of mock-treated Arabidopsis roots from the BFA dataset with the vacuole marker profile indicated in dark blue and the fractionation patterns of AP1/2B1 and AP1/2B2 annotated. (D) Part of the UMAP of the BFA treatment with the AP2 subunits annotated. (E) Part of the UMAP of the BFA treatment with the AP1 subunits annotated. (F) UMAP of the comparison of the *gnom^fwr^*mutant to wildtype seedlings with the AP4 subunits annotated. (G) Distribution patterns of the AP4 subunits in Arabidopsis seedlings with the chloroplast indicated in pink. (H) Distribution patterns of the AP4 subunits in Marchantia thallus with the proteasome indicated in green.

As a user case, we explored the behavior of Adaptor Protein (AP) complexes; obligate complexes that regulate of protein trafficking through facilitating vesicle formation. The 5 family members all display unique subcellular localizations^89–91^. We plotted the AP1 and AP2 subunits, including the shared AP1/2 subunits, to the BFA- and mock-treated root map (Figure 7A). This showed that AP1 localizes to the cytosol, while AP2 localizes to the vacuole and the shared AP1/2B1/2 subunits localize closer to AP2 (Figure 7A). The fractionation profiles of AP2 in the mock-treated root confirm that the AP1/2B1 and AP1/2B2 subunits behave similarly to the AP2 subunits (Figure 7B, C). Treatment with BFA affected the AP complexes differently: the AP2A1 and AP2M (differential localization score 0.998 and 0.512) subunits are differentially localized, while for AP1 no subunits are affected (Figure 7D, E). For AP4, only the AP4B and AP4E subunits are affected in the *gnom^fwr^* mutant (differential localization scores 1 and 0.992) (Figure 7F). Furthermore, the fractionation patterns for AP4 in the Arabidopsis seedling are significantly different compared to the Marchantia thallus (Figure 7F, G), suggesting neo-functionalization of the complex, consistent with the enrichment of GO-terms for protein trafficking related proteins in proteins with large Evolutionary Distance score between the two species (Figure 5M). Thus, navigating sets of proteins across these spatial proteome maps allows the formulation of new hypotheses regarding localization, function, conservation and diversity.

## Discussion

We present an optimized protocol to globally map protein subcellular localizations in plants and generated a number of datasets of subcellular proteomes in the flowering plant *Arabidopsis thaliana* and the liverwort *Marchantia polymorpha*. Each includes thousands of proteins for which a high-confidence subcellular localization can be inferred. The annotations are supported by cross-referencing using confocal microscopy *in vivo,* showing high accuracy in Arabidopsis roots and slightly lower accuracy in Marchantia. We also show that integration with complementary datasets offers a new layer of information. We demonstrate an evolutionary comparison of proteome localization and conclude that the plant subcellular proteome is deeply conserved.

To our knowledge, there is no resource on the plant subcellular proteome with this depth and resolution published to date. In generating validated maps, we increase the number of proteins whose subcellular localization is experimentally tested by several-fold in Arabidopsis, more than an order of magnitude in Marchantia. When we compare our data to recent publications that also use label-free DIA to obtain subcellular spatial proteomes from eukaryotic cell cultures, the depth and resolution are similar or higher^92,93^. Importantly, this method can be performed on wild-type plants, eliminating potential localization artefacts caused by protein tagging and enabling subcellular proteome mapping in plant species that currently cannot be genetically modified. Additionally, while the maps generated here require extensive collection of fractions and replication, the reported fractionation patterns can be leveraged to design simpler experiments that require less separation of certain subcellular localizations. Since we show high conservation of the plant subcellular proteome, the Arabidopsis fractionation patterns can guide the annotation of data in less-studied plant species by comparing proteins from the same orthogroups. Lastly, proteins with a high evolutionary distance score in the comparison between Marchantia and Arabidopsis provide interesting candidates to study protein neo-functionalization and GO-term analysis of these proteins strongly suggests that protein trafficking has been specifically altered during evolution.

In our initial maps, there is a key step that assigns identities to statistically defined clusters, based on the distribution of well-curated and well-established markers for organelles. It should be clear that, as the number of such markers increases, the re-annotation of the existing maps will allow the refinement of the subcellular spatial proteome into more granulated patterns. As an example, sub-clusters can be found in nearly all organellar clusters. Given that the initial clustering is a direct reflection of the sedimentation behavior of each protein, it is likely that even small sub-clusters report on biologically relevant structures. We used a relatively conservative approach where only clusters that were identified by multiple markers that were unambiguous in localization were assigned an identity. We expect that frequent updating of these maps with novel marker sets will increase the utility of the maps. Furthermore, the increasing recognition of membrane-less organelles and large protein superstructures (such as condensates) as biologically relevant entities^94–98^, and the preponderance of LLPS-prone proteins to localize in “ribosome” and “unknown” identity clusters, bodes well for the use of these maps in dissecting protein localization beyond “classical” organelles.

We also show that datasets from two conditions can be successfully compared to identify translocating proteins, which could offer more insights into biological processes. We show that we can identify both known and undescribed responses to the widely used drug BFA using comparative subcellular proteomics, which is supported by confocal microscopy. Additionally, we show that the translocating proteins that we identify are robust and likely biologically relevant, by performing comparative subcellular proteomics on the weak *gnom^fwr^* allele, a known BFA target, which indeed showed a significant overlap in the identified translocating proteins. While these dynamic maps were prepared as a proof-of-concept, we recognize that their further exploration can be a powerful starting point to better understand the mechanisms underlying membrane trafficking and their cargoes, as we demonstrated by some examples from the family of AP-complexes.

In addition to obtaining better resolution, we see a range of possibilities to capitalize on and expand plant subcellular proteomics. Clearly, the expansion to a broader range of plant species – including ones with unique or specialized organelles (e.g. pyrenoids, oil glands, centrioles) will assist in building an experimentally grounded framework for protein-based evolutionary cell biology. Complementary approaches such as comparative Expansion Microscopy^99^, comparative phosphoproteome profiling^53^, and comparative protein interaction profiling^100^ can be integrated with spatial proteome maps to formulate new hypotheses and guide the path to new biology. Using carefully selected treatments and mutants, it will be possible to generate more insights into a broad variety of processes, such as protein trafficking networks, biotic and abiotic stresses, and cell signaling. Additionally, the protocol could also be adapted to be applied to plant cell cultures, or it could be optimized to generate spatial maps of post-translationally modified proteins, as well as chemically cross-linked proteins.

### Limitations of the study

While the maps provided here increased the number of proteins with an experimentally grounded subcellular localization in plants by a large factor, the MS-based method is inherently limited to only those proteins that can be directly detected. Regulatory proteins are often low in abundance and are likely underrepresented in these datasets. However, mass spectrometers and the downstream processing are continuously becoming more sensitive, as we demonstrated by comparing two mass spectrometers. Hence, this limitation will be reduced over time. Furthermore, in comparing mutants with wild-type, it is possible that mutant tissue integrity (e.g. cell-cell adhesion, cell wall stiffness, membrane properties) affects lysis. Such effects may complicate direct comparisons to the wildtype, as one cannot assume that liberation of cell contents is fully comparable. Furthermore, the quality of cluster annotations is dependent on the presence of well-established marker proteins, which are generally scarce in plants. We expect that resources such as this one will in fact, lead to the identification of more markers and thus more compartments. Lastly, even the simplest tissue profiled here (roots) consists of multiple cell types that differ in composition, size, and likely in subcellular organization. Hence, the present maps average such cellular heterogeneities and likely present a simplified, unified picture of subcellular organization. While it will remain challenging to isolate individual cell types without a destructive process that nullifies cellular organization (e.g. cell wall digestion and protoplast sorting), the problem can partly be resolved by adapting the protocol for use on uniform plant cell cultures.

## Supporting information

Oligonucleotides

## Acknowledgements

We thank our team members for continuous support and helpful discussions, Sjef Boeren for advice on mass spectrometry, Kathryn Lilley for suggestions, Lisa Breckels for providing the code for the Shiny apps, Alexis Maizel for sharing a published plasmid, Enrico Scarpella, Yasin Dagdas, Juan Dong, Michael Sauer, Takashi Ueda, Kentaro Tamura, Andre Kuhn, and Enrique Rojo for sharing published lines, and Anna Jonkers, Cor Wolfs, and Simon Lindhoud for experimental support. This work was supported by a PhD fellowship from the Graduate School Experimental Plant Sciences (Wageningen University) to M.v.S., by the European Research Council (ERC AdG “DIRNDL; contract 833867) to D.W., the Sector Plan Biology of the Dutch Ministry of Education, Culture and Science, and the Netherlands Organization for Scientific Research (NWO; CropXR) to M.R..

## Declaration of interests

The authors declare no competing interests.

## Materials and methods

### Key Resources table

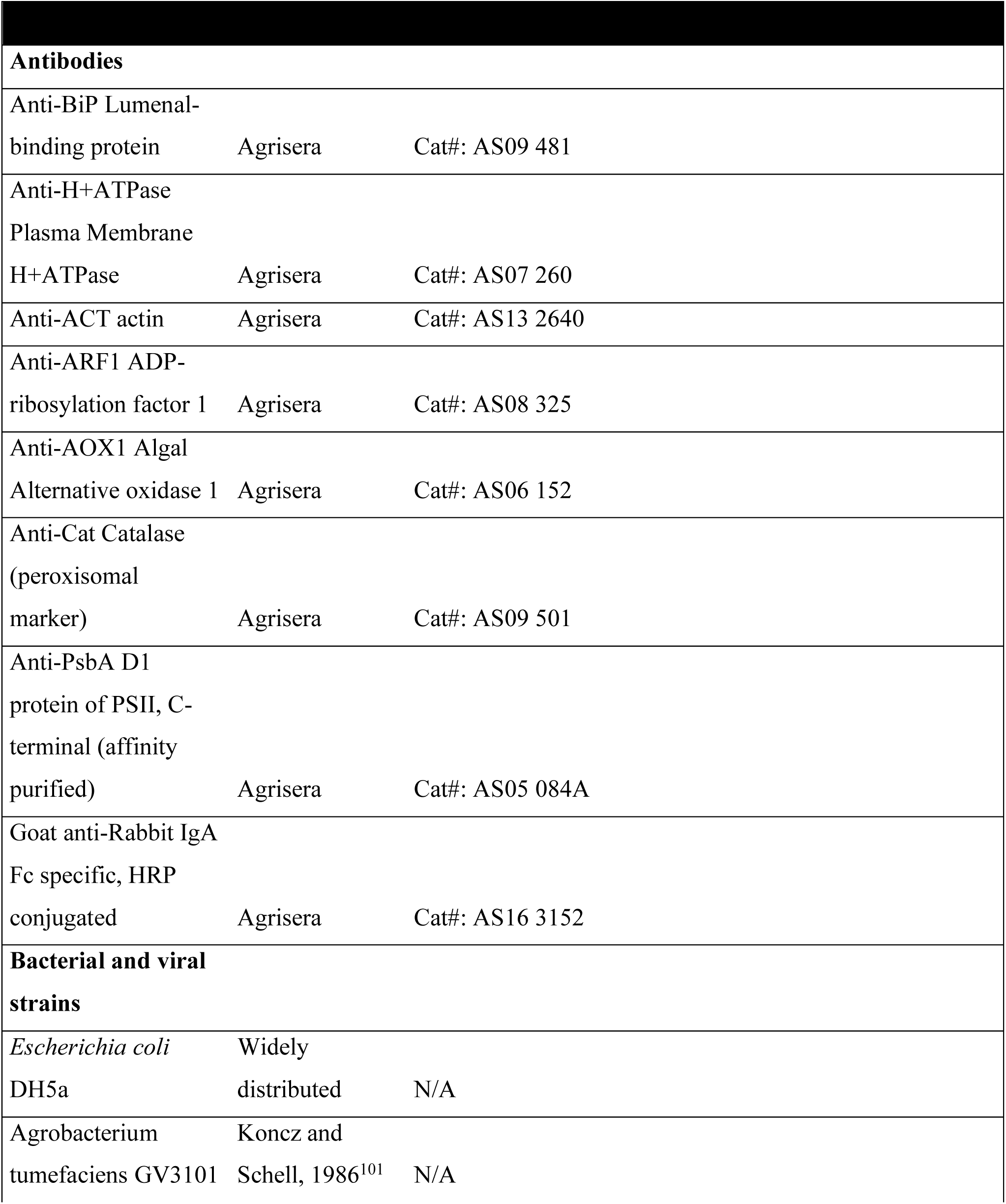
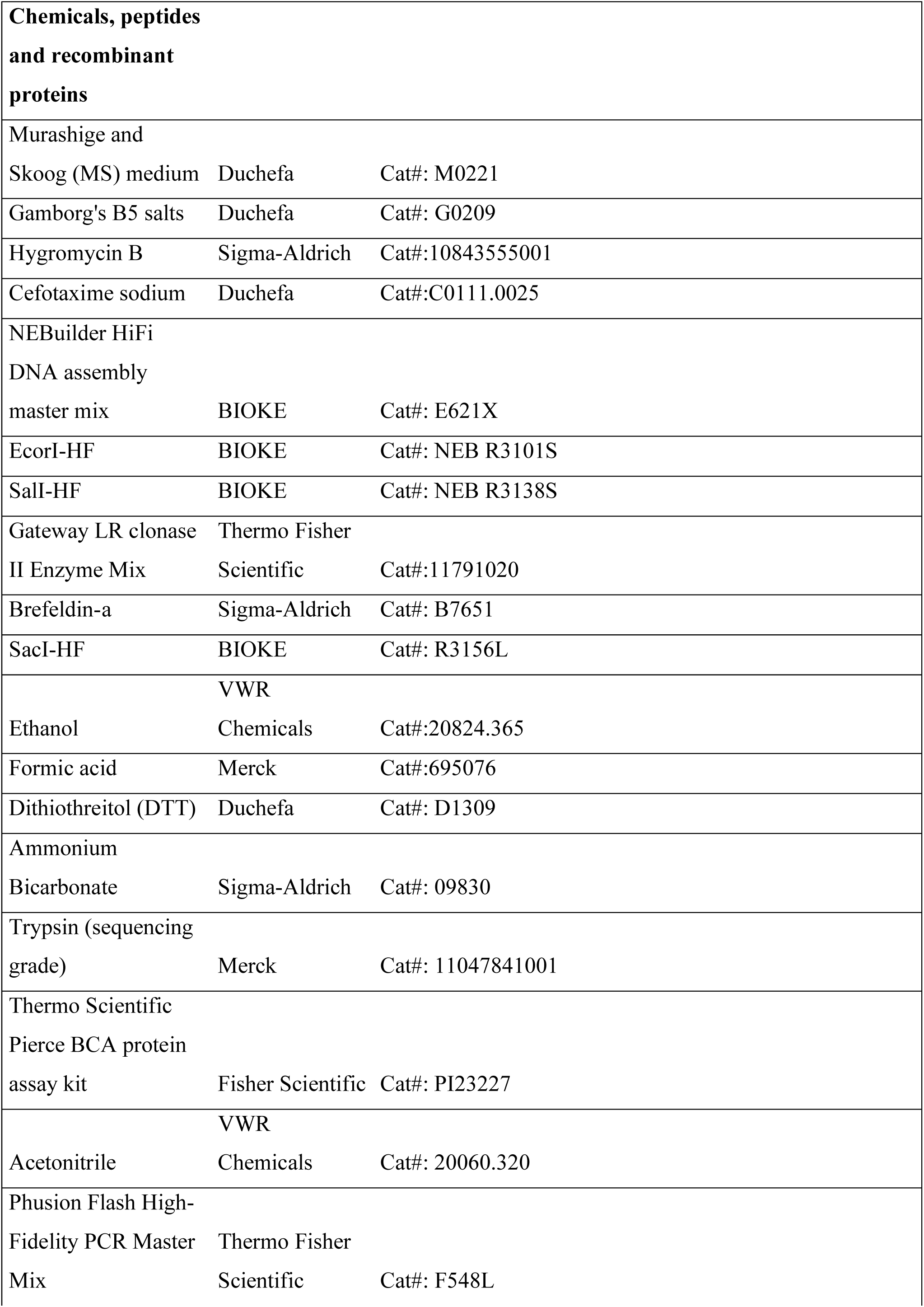
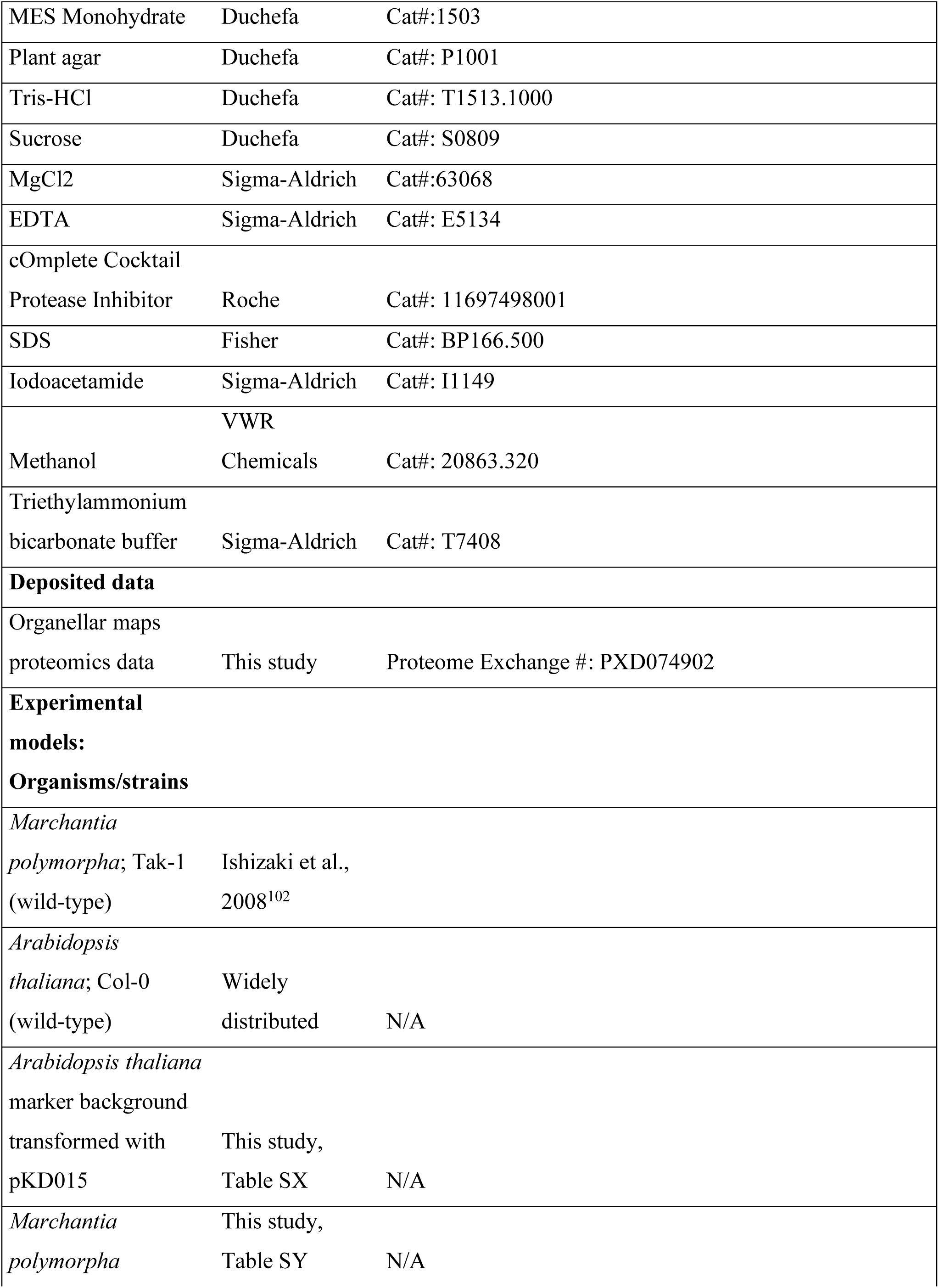
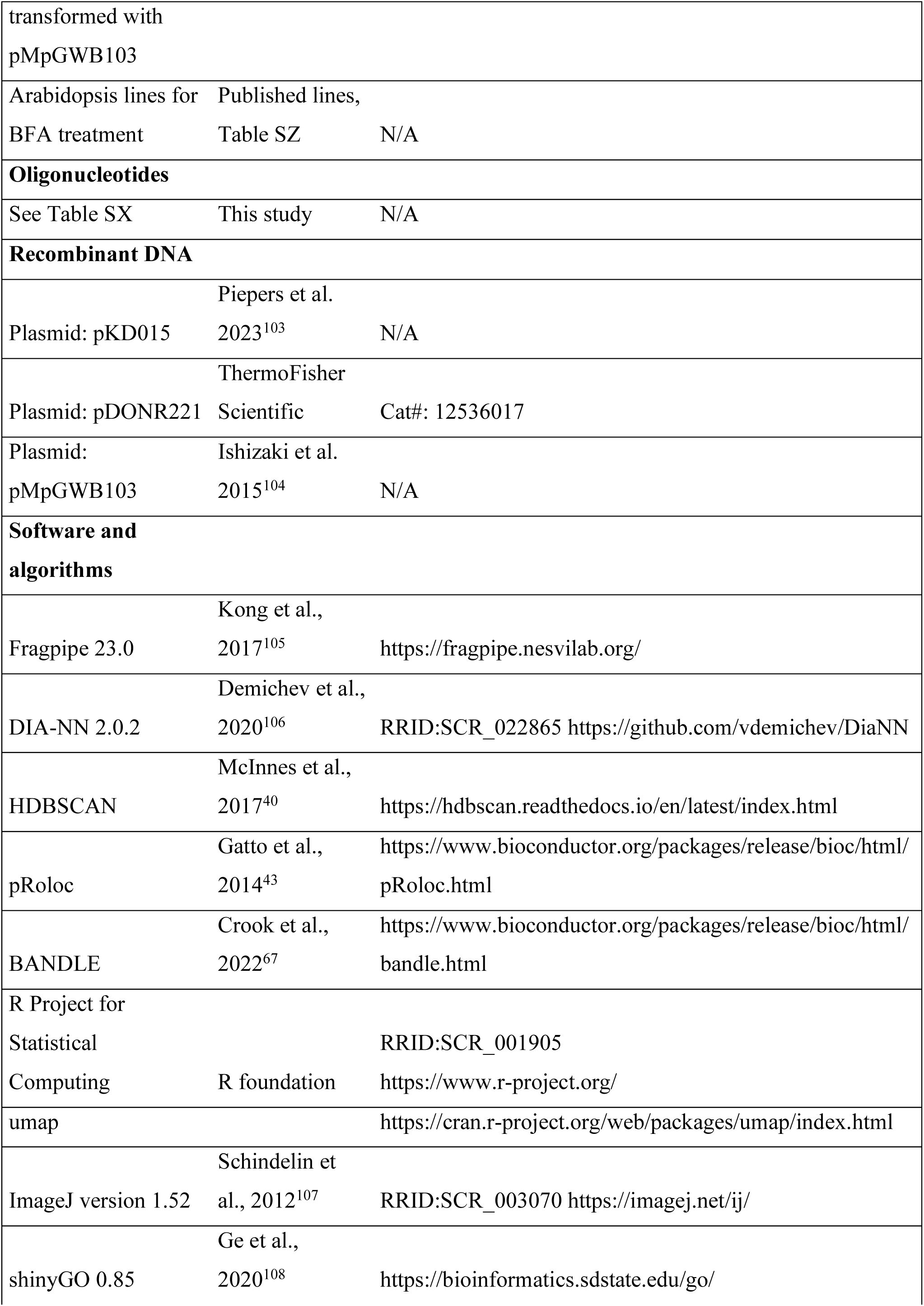
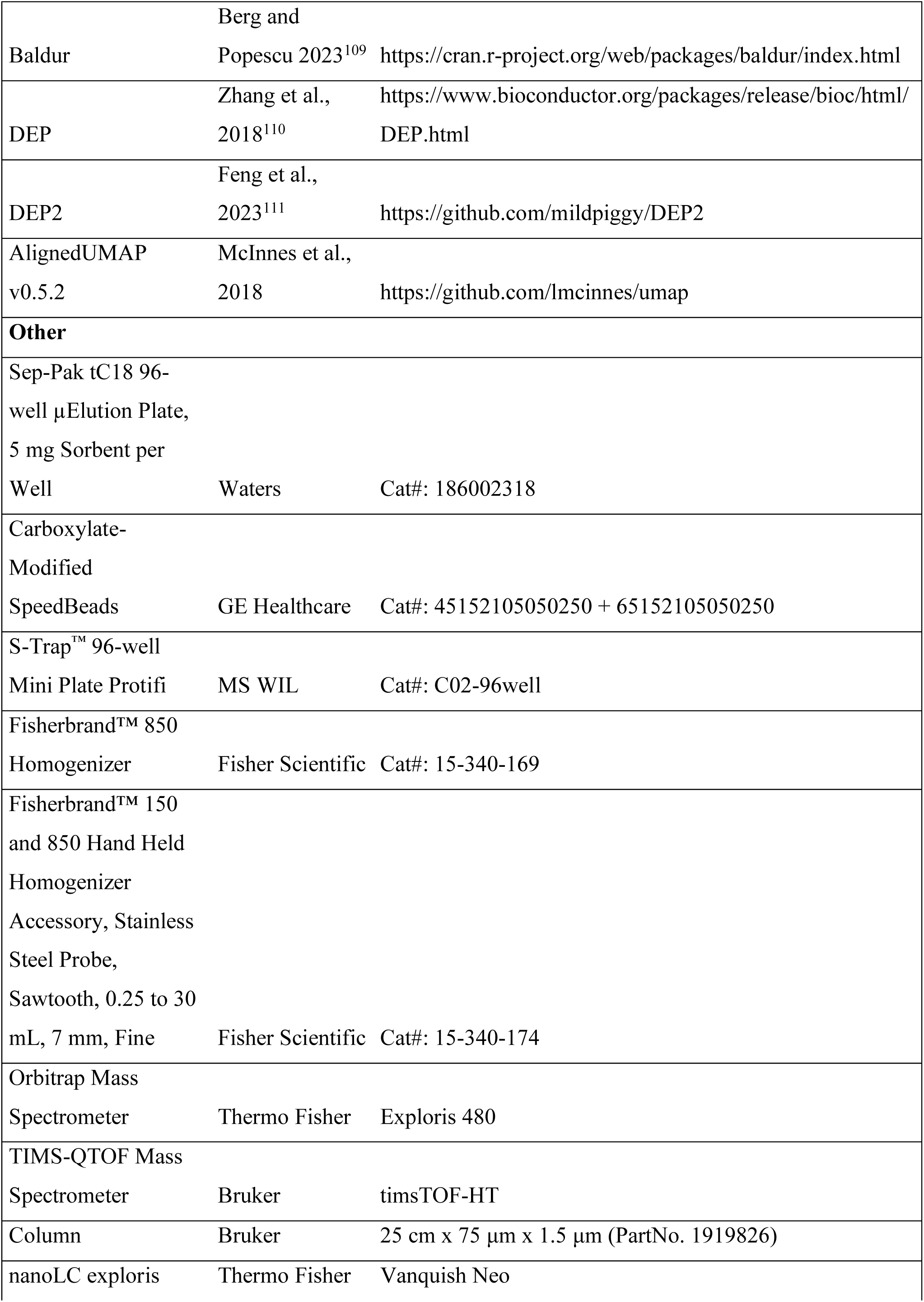
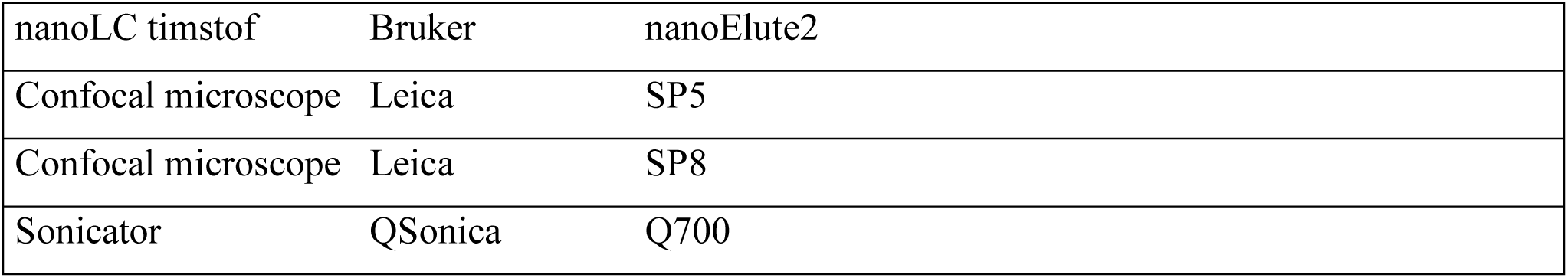

### Resource availability

All new genetic materials generated in this study are available from the lead author upon request, without restrictions. All proteome data has been deposited in ProteomeXchange (PXD074902), and can be accessed through Shiny apps^88^: [Roots](https://weijerslab.shinyapps.io/spatial_proteomics_roots_arabidopsis_thaliana/), [Seedling](https://weijerslab.shinyapps.io/spatial_proteomics_seedling_arabidopsis_thaliana/), [Marchantia](https://weijerslab.shinyapps.io/spatial_proteomics_marchantia_polymorpha/), [BFA treatment](https://weijerslab.shinyapps.io/spatial_proteomics_dynamic_bfa_treatment/), [gn-fwr](https://weijerslab.shinyapps.io/spatial_proteomics_dynamic_gnfwr/).

## Materials

### Plant materials and culture conditions

*Arabidopsis thaliana*, ecotype Colombia-0 (Col-0), including the *gn^fwr^* mutant and transgenic lines, were grown vertically on half-strength Murashige and Skoog basal medium^112^ at pH 5.7, supplemented with 0.8% agar for 7 days. The *gn^fwr^* allele was first described by Okumura et al^37^. For the validation of the Arabidopsis root data *in vivo*, the WAVE lines WAVE-1R, WAVE-6R, WAVE-9R, WAVE-13R, WAVE-22R and WAVE-138R^44^ (NASC stock facility), and mit-GFP^45^ were used. (Table S1). For the validation of the BFA treatment, the MTV1^65,69^, VPS3^113^, IMPA2, IMPA4^70^, IYO^72^, RAF20^53^, BSL1^114^ lines were used (Table S3).

*Marchantia polymorpha* Takaragaike-1 (Tak-1) was used as wild-type. The plants were cultured on ½ strength Gamborg B5 medium at 22°C with constant white light (40 μmol photons m–2s –1). Transformation of *Marchantia polymorpha* spores was performed as previously described^115^. For transformants selection, the B5 plates were supplemented with 10 mg/L hygromycin and 100 μg/ml cefotaxime.

All the plants were grown under 16 hours of light and 8 hours dark at 75% humidity, at a light intensity of 90-100 µmol photons m^-2^s^-1^.

## Methods

### Cloning and transformations

For Arabidopsis transformations, the pKD015 expression vector^103^ was used to make C-terminal translational mTurq2 fusions driven from the *UBQ10* promotor. The vector was digested using EcorI and SalI. Amplicons were obtained by PCR from genomic DNA using primers with vector specific overhangs (Table S4). Ligation was performed using HiFi (NEB) and confirmed by Sanger Sequencing. Constructs were stably expressed in Arabidopsis by floral dip^116^. In Arabidopsis, respective marker background plants were used (Table S1).

For Marchantia transformations, protein TurboID-YFP fusion were generated. Genomic fragments of the genes of interest without stop codon were amplified from Marchantia genomic DNA (Table S4). Gateway entry vectors were obtained by transferring PCR-amplified sequences into a linearized pDONR221 (amplified with GAAGCCTGCTTTTTTGTACAAAGTTGGC and GACCCAGCTTTCTTGTACAAAGTTGGC) via NEBuilder® HiFi DNA Assembly (New England Biolabs). The resulting entry vectors were transferred into a modified destination plasmid pMpGWB^104^ by LR reaction using Gateway® LR Clonase® II Enzyme Mix (Thermo Fisher Scientific). The modified destination vectors carry the TurboID-YFP cassette cloned in the pMpGWB103^104^ via SacI restriction site for C-terminal fusion. Transformation of Tak1 spores was performed as previously described (Kato et al, 2015) using *Agrobacterium* strain GV3101:pMp90^101^. Oligonucleotides used in that study are listed in Table S4, the obtained plant lines are listed in Table S2.

### Imaging and image analysis

For the validation of the predictions in Arabidopsis roots, all selected proteins were expressed as fusions to mTurquoise2, which was excited at 442 nm and detected between 450 and 550 nm. Most background lines carry mCherry, which was excited at 561 nm and detected at 570 to 670 nm. The mit-GFP background was excited at 488 nm and detected at 550 to 650 nm to avoid bleed through of the mTurquoise2 signal. In this background, the detection window of mTurquoise2 was reduced to 450-480 nm. All images were obtained using a Leica SP5 confocal microscope fitted with hybrid detectors and an Argon laser. Raw images were analyzed using ImageJ^107^, where the LUT was set to green for the channel with the protein of interest and magenta for the background. The brightness and contrast of the channels were adjusted individually to optimize the comparison of the channels.

For the validation of predictions in Marchantia, all microscopy was performed on gemmae using a Leica SP5 confocal microscope fitted with hybrid detectors and an Argon laser. The YFP fluorophore was excited using 514 nm laser line at 20% power. Fluorescence was detected between 520-550 nm with hybrid detectors set to photon counting mode. Images were analyzed using ImageJ^107^.

Arabidopsis seedlings were treated with 10 µM BFA on microscopy slides in liquid half strength MS medium^112^ or with equal volumes ethanol in half strength MS medium for 1 hour. Lines with GFP were excited at 488 nm and detected with 495 to 550 nm, and lines with YFP were excited at 512 nm and detected at 520 to 575 nm. All images were obtained using an SP5 or SP8 confocal microscope (Leica). Images were analyzed using ImageJ^107^. For every line, brightness and contrast were set at the same values and the Fire LUT was used.

### Spatial proteomics sample preparation

Harvesting of plant materials was performed using fresh tissue, without snap-freezing. For the BFA treatment, the plates with 7-day-old seedlings were placed horizontally and flooded with 10 µM BFA dissolved in ethanol diluted in half strength liquid MS medium or an equal volume ethanol diluted in liquid half strength MS medium for the control, and after draining excess fluid, incubated for 30 minutes. After the treatment, roots were harvested using a scalpel. Arabidopsis seedlings, roots and Marchantia thallus were harvested from the plates directly without treatment, the Arabidopsis roots were harvested with a scalpel as described for the BFA treatment. Samples that are directly compared (treatment vs mock or mutant vs wild type) were collected pairwise on the same day.

Plant tissues were harvested into pre-cooled lysis buffer (25 mM Tris-HCl, pH 7.5, 250 mM Sucrose, 0.5 mM MgCl2, 0.2 mM EDTA,1mM DTT, 1x Protease Inhibitor Cocktail) that was placed on ice. For Arabidopsis roots, 15-20 plates were used for each replicate, for Arabidopsis seedlings and Marchantia thallus ∼1 plate was used per replicate, all densely sown on a layer of nylon mesh (100 µm mesh size). The tissues were homogenized using an automated homogenizer with a 7 mm probe (ThermoFisher). For Arabidopsis roots 5x 5 second pulses at 15000 RPM were used, and for the Arabidopsis seedlings and Marchantia thallus 4x5 second pulses at 25000 RPM were used. During the pulses, the samples were moved vertically to ensure that all tissue was homogenized. Subsequently, the samples were transferred to a Dounce homogenizer and homogenized using 15 strokes. After passing through a 100 µm mesh filter, samples were ready for centrifugation.

To remove unlysed cells, samples were centrifuged at 200xg for 5 minutes. Subsequently, the supernatant was collected, and organelles were separated at increasing speeds. After every centrifugation step, the pellet was kept on ice and the supernatant transferred to a new tube.

When using the LOPIT-DC method^38^ the centrifugation steps were 1000xg 20 minutes (1K pellet), 3000xg 10 minutes (3K), 5000xg 10 minutes (5K), 9000xg 15 minutes (9K), 12000xg 15 minutes (12K), 15000xg 15 minutes (15K), 30000xg 20 minutes (30K), 79000xg 45 minutes (79K) and 120000xg 45 minutes (120K) and the supernatant was kept as the last cytosol fraction (cyt), yielding 10 fractions in total.

When using the DOMs method^8^ the centrifugation steps were 1000xg 20 minutes (1K pellet), 3000xg 10 minutes (3K), 5400xg 15 minutes (5K), 12200xg 20 minutes (12K), 24000xg 20 minutes (24K), 78400xg 30 minutes (78K) and the supernatant was kept as the cytosol (cyt) fraction, yielding 7 fractions in total. Additionally, whole proteome samples (wp) were taken after passage through the mesh filter at the last step of tissue lysis, yielding 11 samples per repetition for LOPIT-DC and 8 samples for DOMs, respectively.

The pellets were resuspended in SDS lysis buffer (2% SDS, 50 mM Tris-HCl, pH 8.1) and the cytosolic fraction and whole proteome samples brought to the same SDS concentration by adding 5x SDS lysis buffer (10% SDS, 50 mM Tris-HCl, pH 8.1). The resuspended samples were heated at 72 °C for 5 minutes and sonicated (Qsonica) for 15 30-second on/off cycles at 70% intensity. Protein concentrations were measured in triplicate using a BCA assay (ThermoFisher).

### Western Blots

For the initial testing of the protocol, Western Blots with antibodies for cellular compartments were used. To start, 10 µg of protein for each fraction was added to SDS loading dye and MQ water was added to obtain 1/4^th^ strength of the SDS loading dye in the final samples. After heating for 10 minutes at 95°C, samples were loaded to a pre-cast gel (Invitrogen) and ran in MOPS SDS running buffer (Invitrogen) and ran at 90-120 Volt for 1-1.5 hours. Proteins were transferred to a nitrocellulose membrane using a Power Blotter (Invitrogen). After the transfer, membranes were incubated in blocking buffer (5% BSA in TBST) for 45 minutes. Next, membranes were incubated with the manufacturer recommended concentration of primary antibody dissolved in blocking buffer overnight. Membranes were washed 3 times for 5 minutes with TBST before applying the secondary antibody at the manufacturer recommended concentration for 45 minutes. Afterwards, membranes were washed 3 times for 5 minutes with TBST before visualization. Visualization of bands was done using the Amersham™ ECL Detection Reagents (Cytiva) and imaging performed using a ChemiDoc imager (Bio-Rad).

### Protein Aggregation Capture (PAC)

Calculations were made to obtain 10 µg protein input for every fraction and protein concentrations were normalized using SDS lysis buffer. Proteins were subsequently acetylated using iodoacetamide 20 mM for 30 minutes in the dark. The PAC beads (GE Healthcare) were washed twice with 1 ml mQ, before incubation with the samples. Beads were used in a 1:10 protein: bead ratio and were mixed well by pipetting with 70% ACN and the samples, which were incubated for at least 20 minutes at room temperature. Afterwards, 3 washes were performed with 100% ACN and 2 washes with 70% ethanol. Trypsin was added in 50mM ABC in a 1:100 ratio to the protein concentration and digestion was performed overnight.

### S-Trap

The *gn^fwr^* mutant and the corresponding control samples were prepared using S-Trap instead of PAC. When sample prep was performed using S-Trap, the pellets were resuspended in SDS resuspension buffer (5% SDS in 50 mM TEAB, pH 7.55). The BCA assay was performed in the same manner as described above. There was a protein input used of 10 µg per sample for every fraction, which were normalized to the same concentration using SDS resuspension buffer. Proteins were subsequently acetylated using iodoacetamide 20 mM for 30 minutes in the dark. Afterwards, samples were acidified to a final concentration of 1% FA.

Before loading onto the S-Trap plate, samples were mixed well with 6x the volume binding buffer (90% methanol in water, 100 mM TEAB, pH 7.1). The samples were loaded by centrifuging at 1500 xg for 2 minutes. Afterwards, the plates were washed three times with 200 µl binding buffer per well by centrifuging at 1500 xg for 2 minutes. To digest the proteins, 100 µl 50 mM ABC with trypsin in a 1:100 ratio the protein was added. The plates were covered with a lid and sealed with parafilm to avoid evaporation during the overnight digestion. The next day, the trypsin solution was centrifuged into a clean receiver plate for 2 minutes at 1500 xg. Afterwards, peptides were eluted in the same receiver plate using 120 µl 80% ACN in 0.1% FA. Samples were reduced to about 20% of the original volume by vacuum centrifuging. Both samples from S-Trap and PAC were subsequently desalted using C18 clean-up.

### C18 clean-up

The digested samples were acidified to a pH of 2-3 using 10% FA and cleaned-up using Sep-Pak C18 96-well plates (Waters). The plate was activated by one wash with 80%ACN/0.1% FA and two washes with 0.1%FA at 1500xg for 2 minutes using a swing-bucket centrifuge. The samples were loaded for 10 minutes at 750xg. Samples were washed twice with 0.1% FA before being eluted using 80%ACN/0.1% FA, all for 2 minutes at 1500xg. Samples were dried in a vacuum centrifuge at 45 degrees for 45 minutes. Lastly, the peptides were resuspended in 0.1% FA.

### MS measurements

Mass spectrometry on fractionated samples was essentially performed as described previously^53^ with the exception that data independent acquisition was performed. For the *gnom^fwr^* data 2 µl samples were loaded on a Pepsep advanced 25 cm x 75 µm x 1,5 µm (Bruker) at a constant pressure of 800 bar with 1ml/l HCOOH in water and eluted at a flow of 0.3µl/min with a 28 min linear gradient from 2% to 35% acetonitrile in water with 1 ml/l HCOOH with a nanoElute2 (Bruker). Data was acquired using an pydiAID optimized synchroPASEF acquisition scheme, 125m/z with 4slices^117^ with a cycle time of 0.53 seconds. TIMS ramp was set to 100ms. The collision energy was decreased as a function of the IM from 59 eV at 1/*K*_0_ = 1.6 V cm^−2^ to 20 eV at 1/*K*_0_ = 0.6 V cm^−2^, and the IM dimension was calibrated with three Agilent ESI Tuning Mix ions (*m/z*, 1/*K*_0_: 622.02, 0.98 V cm^−2^, 922.01, 1.19 V cm^−2^, 1221.99, and 1.38 V cm^−2^).

### Raw data processing and quantification

For DDA acquired data Thermo RAW files were analyzed with MSFragger^105^ version 23 using the built in “LFQ-MBR” workflow. For DIA acquired data Thermo RAW files or Bruker .d files were analyzed using DIA-NN^106^ version 2.0.2. Spectral libraries were predicted from Arabidopsis Thaliana (UP000006548) or Marchantia polymorpha proteomes (UP000077202) retrieved from UniProt. MSFragger combined_ion.tsv or DIA-NN report.parquet files were used as input for quantification using DirectLFQ^118^ with an in house generated Python script. Data was filtered on protein and peptide q values ≤0.01 and a minimum number of ion intensities of 2. The mass spectrometry proteomics data have been deposited to the ProteomeXchange Consortium via the PRIDE^119^ partner repository with the dataset identifier PXD074902.

### Computational methodology for spatial proteome characterisation

Data analysis was performed using R^120^ packages pRoloc^43^, BANDLE^67^, UMAP^39^, DEP2^111^, Baldur^109^. Raw data was filtered on protein groups identified with at least 2 peptides. DirectLFQ intensities values were feature scaled using row min-max normalization. Normalized data was used as input for the HDBSCAN algorithm as implemented in Python using a min cluster size of 20 and min samples of 80. Proteins belonging to the identified clusters were searched using SUBA5^41^ and experimental data reported in the UniProt database^42^ to build a marker set. Markers were visualized on a UMAP generated using n_neighbors of 20, min_dist of 0.1 and Euclidean distance as metric. Pairwise marker correlations were calculated using the matrixTest package in R.

Predictions of localizations was made with a final marker set using a T-Augmented Gaussian Mixture model Markov-chain Monte-Carlo (TAGM-MCMC)^121^ run with 20000 iterations, burn-in of 10000, thinning of 20 and 6 chains. Convergence was assessed using Gelman Rubin diagnostics and non-converged chains were removed. Localisations were reported and visualised with a localisation probability ≥0.99 and an outlier probability ≤ 5e-5. Probabilities not meeting this threshold were reported as unknown.

For differential localisation, the BANDLE package was used implementing a complex full Bayesian framework. Gaussian processes were fit to the marker profiles using default hyperparameters as described in Crook et al. 2022^67^. Next a Dirichlet prior on the mixing weights was set up in such a way that the expected differential localisation is expected to be small and concentrates around a prior probability of 0. TAGM-MCMC was run with 20000 iterations, burn-in of 10000, thinning of 20 and 8 chains. Convergence was assessed using Gelman Rubin diagnostics and converged chains were pooled. Localisations were reported with a BANDLE localisation probability ≥0.99 and a BANDLE outlier probability ≤ 5e-5. Differential localisations were reported with a threshold of ≥0.9999.

For Aligned-UMAP, control and treatment/genotype data was separately processed as described above. Both sets were simultaneously embedded in a lower dimensional space using the following parameters as described by Hein et.al 2025^11^: 20 nearest neighbours, Euclidean distance metric, minimum embedding distance 0.1, 300 epochs, and an alignment regularization factor of 0.002. To map induced translocations following BFA or in the *gnom^fwr^*background, BANDLE calculated differential localisations probabilities with a threshold of ≥0.9999 were mapped on a separate UMAP and were then projected onto the treatment/genotype side of the aligned UMAP through Procrustes analysis as described by Hein.et.al 2025^11^. Evolutionary distance score was calculated using Aligned-UMAP as described above. *Arabidopsis thaliana* seedling data and *Marchantia polymorpha* data was simultaneously embedded in a shared 10-dimensional space. Embeddings were repeated for 200 times to reduce stochasticity. Distance score is defined as the average Euclidean distance of 200 repetitions between *Arabidopsis thaliana* seedling and *Marchantia polymorpha* coordinates. Threshold of distance was defined as 3 times the inter quartile range.

Differential expression analysis was conducted using the DEP/DEP2 packages in R^110^. DirectLFQ calculated intensities were log2 transformed and filtered to have at least 3 valid values in at least one condition. Remaining missing values were imputed using a mixed imputation approach were values missing at random (MAR) were imputed with knn and values missing not at random (MNAR) were imputed based on quantile regression. Differentially expressed proteins were inferred using a Bayesian hierarchical latent gamma mixture regression model (lgmr) as implemented in Baldur^109^. The lgmr model was run with the following parameters: 6 chains, 10000 iterations, warmup of 3000 iterations, adapt delta of 0.95 and maximal tree depth of 11. Chain convergence was checked by plotting all R-hats which should be below 1.1 using Rstan^122^. Significantly differentially expressed proteins were inferred when they passed a probability of error ≤0.05.

### R shiny app

Code for the R shiny apps were kindly provided by Lisa M Breckels^88^. Code was formatted to fit the data files generated in this study.

## Notes

https://weijerslab.shinyapps.io/spatial_proteomics_roots_arabidopsis_thaliana/

https://weijerslab.shinyapps.io/spatial_proteomics_seedling_arabidopsis_thaliana/

https://weijerslab.shinyapps.io/spatial_proteomics_marchantia_polymorpha/

https://weijerslab.shinyapps.io/spatial_proteomics_dynamic_bfa_treatment/

https://weijerslab.shinyapps.io/spatial_proteomics_dynamic_gnfwr/

### Competing Interest Statement

The authors have declared no competing interest.

