## Supplemental Figures and Tables for "High resolution, proteome-wide mapping of subcellular protein localization in plants"

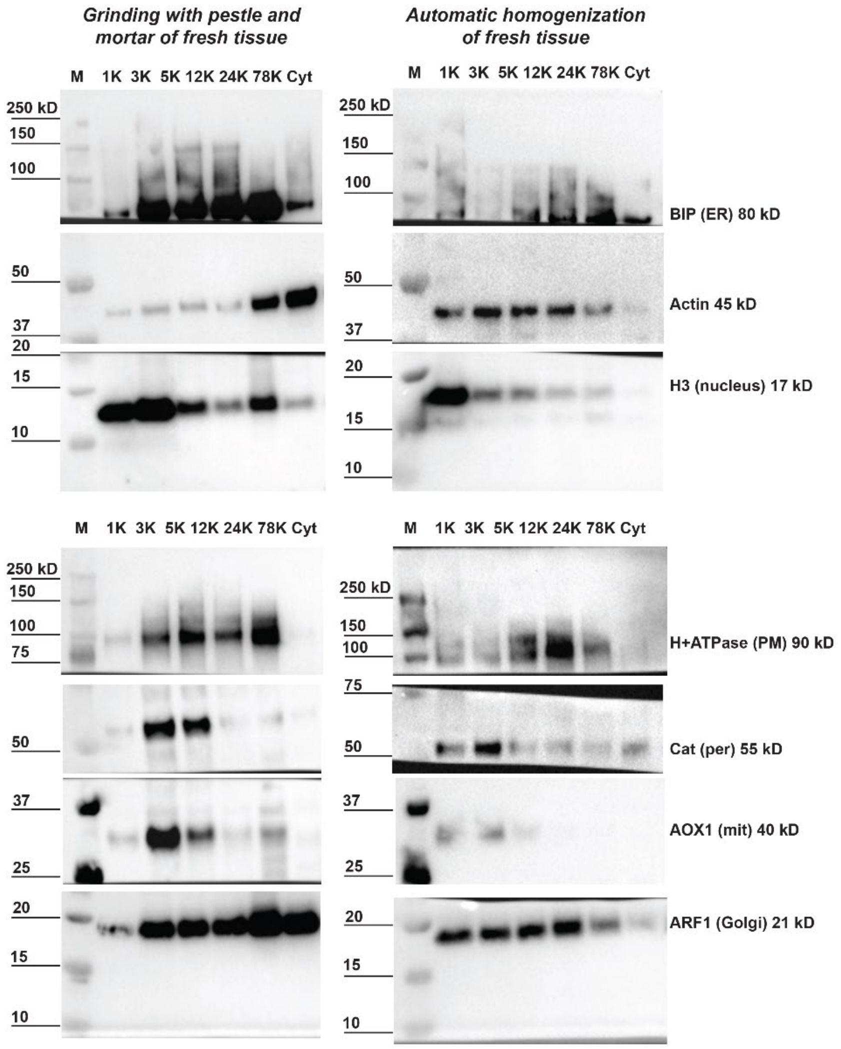


**Figure S1: Optimization of lysis efficiency; related to Figure 1.** Western Blot fractions obtained after lysis of fresh tissue comparing the use of grinding by mortar and pestle and automated homogenization. Primary antibodies for known marker proteins for various subcellular localizations were used and the fractions obtained according to the centrifugation steps described in the DOMS method (details in STAR★Methods).

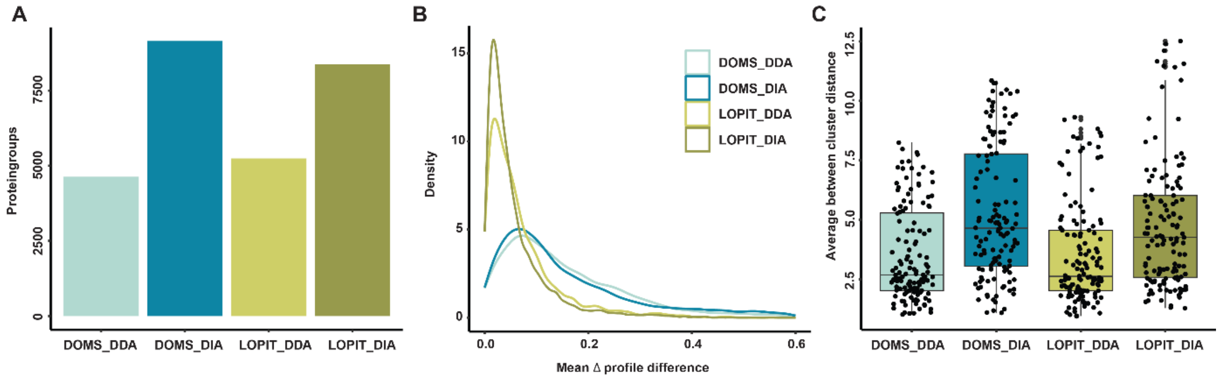


**Figure S2: Comparison of the DOMS and LOPIT-DC methods on Arabidopsis roots, including comparison of measurements in DDA and DIA modes; related to Figure 1.** (A) Number of identified protein groups, the depth increases by approximately 60% when using DIA, both for DOMS and LOPIT-DC. (B) Mean profile differences of fractionation patterns between replicates of the same protein. The LOPIT-DC method measured in DIA mode shows the lowest differences and is therefore the most reproducible. (C) Average between cluster distances and thus fractionation pattern separation is similar when comparing DOMS to LOPIT-DC, both increase when measuring in DIA mode.

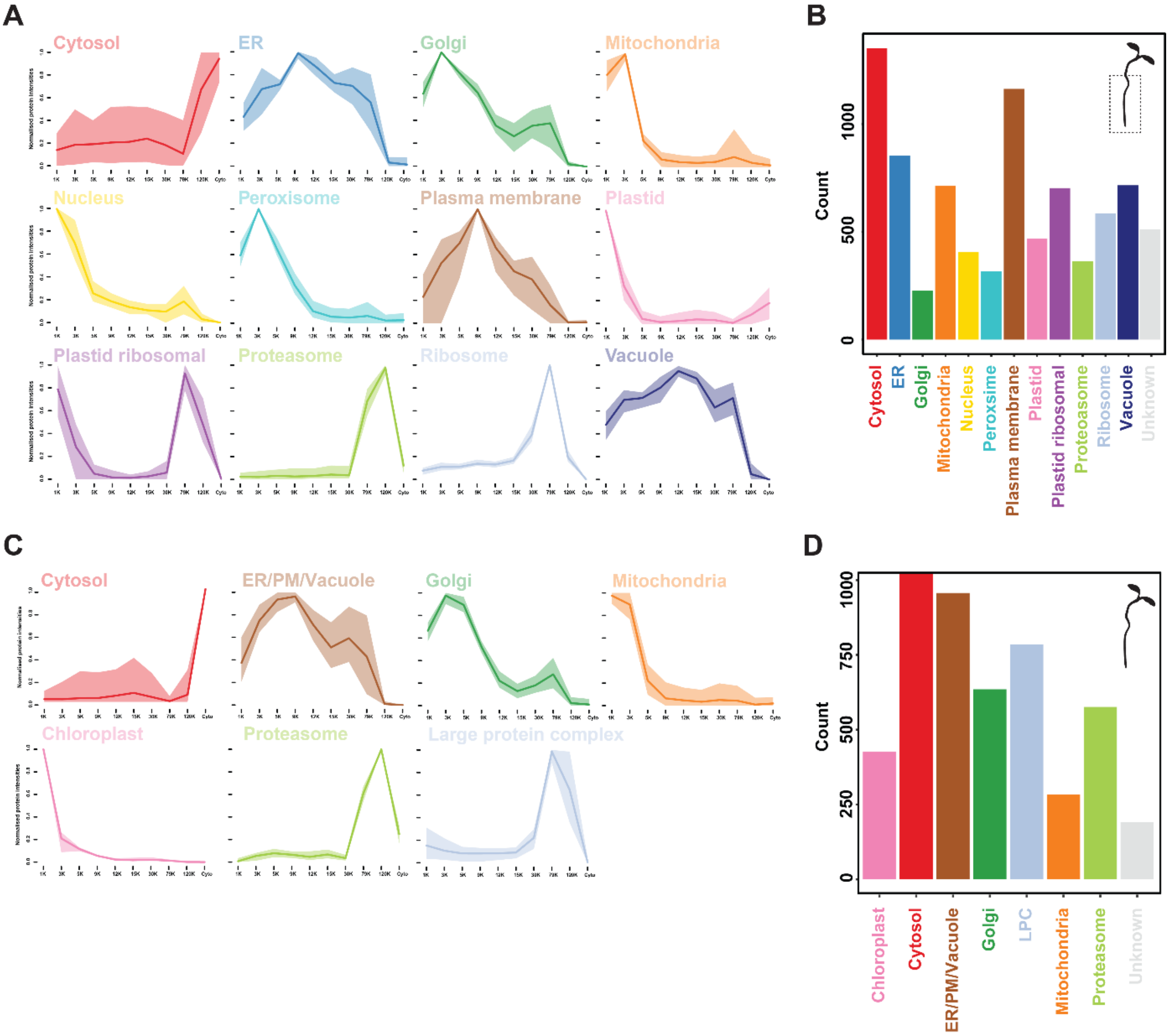


**Figure S3: Obtained fractionation profiles of marker proteins and distribution of proteins over subcellular localizations for Arabidopsis roots and seedlings; related to Figure 1.** (A) Fractionation patterns for the marker proteins for every subcellular localization in the Arabidopsis root dataset. (B) Distribution of the number of annotated proteins for every subcellular localization in the Arabidopsis root dataset. (C) Fractionation patterns for the marker proteins for every subcellular localization in the Arabidopsis seedling dataset. (D) Distribution of the number of annotated proteins for every subcellular localization in the Arabidopsis seedling dataset.

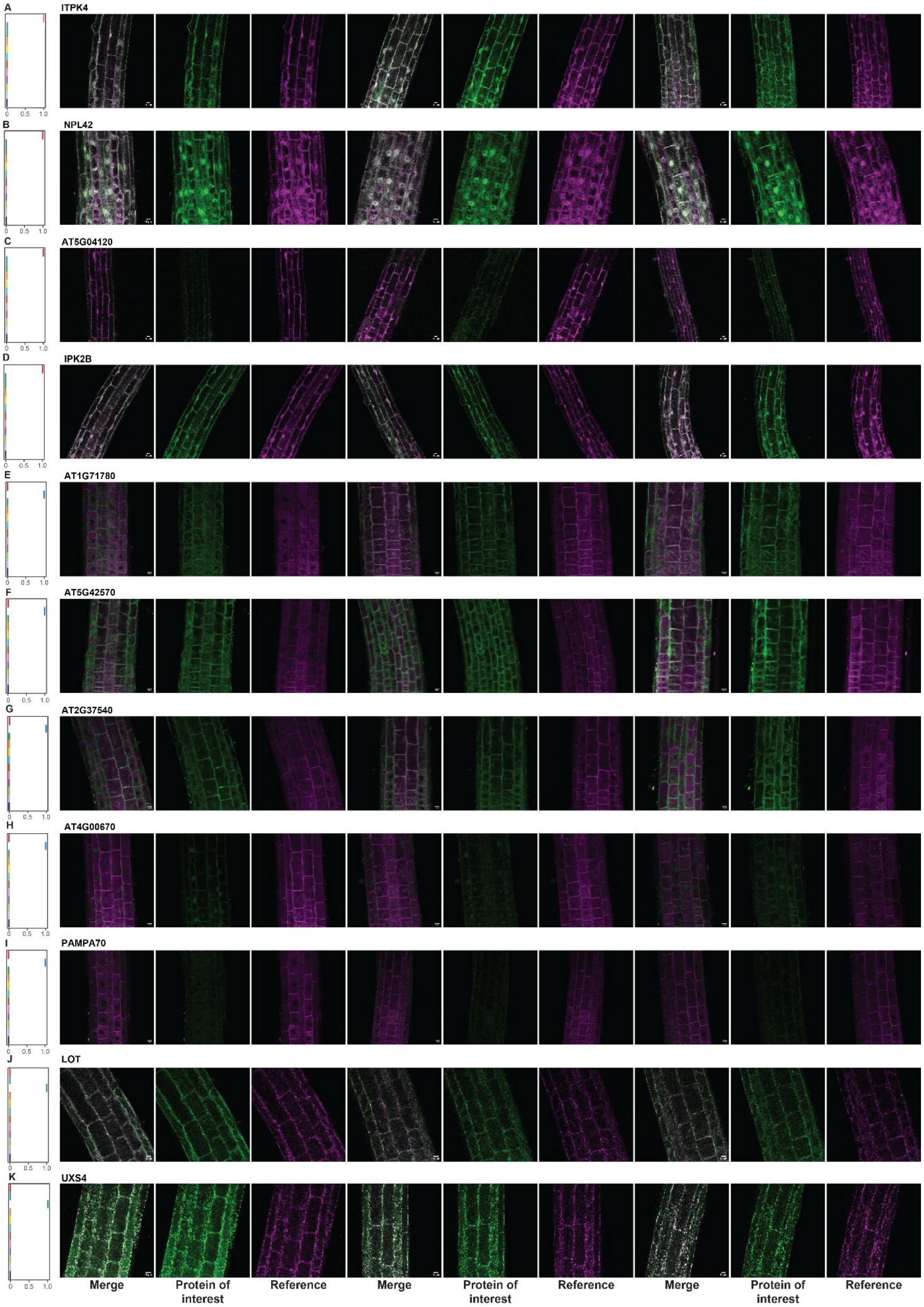
A
ITPK4
B
NPL42
C
AT5G04120
D
IPK2B
E
AT1G71780
F
AT5G42570
G
AT2G37540
H
AT4G00670
I
PAMPA70
J
LOT
K
UXS4
0
0.5
1.0
Merge
Protein of interest
Reference
Merge
Protein of interest
Reference
Merge
Protein of interest
Reference

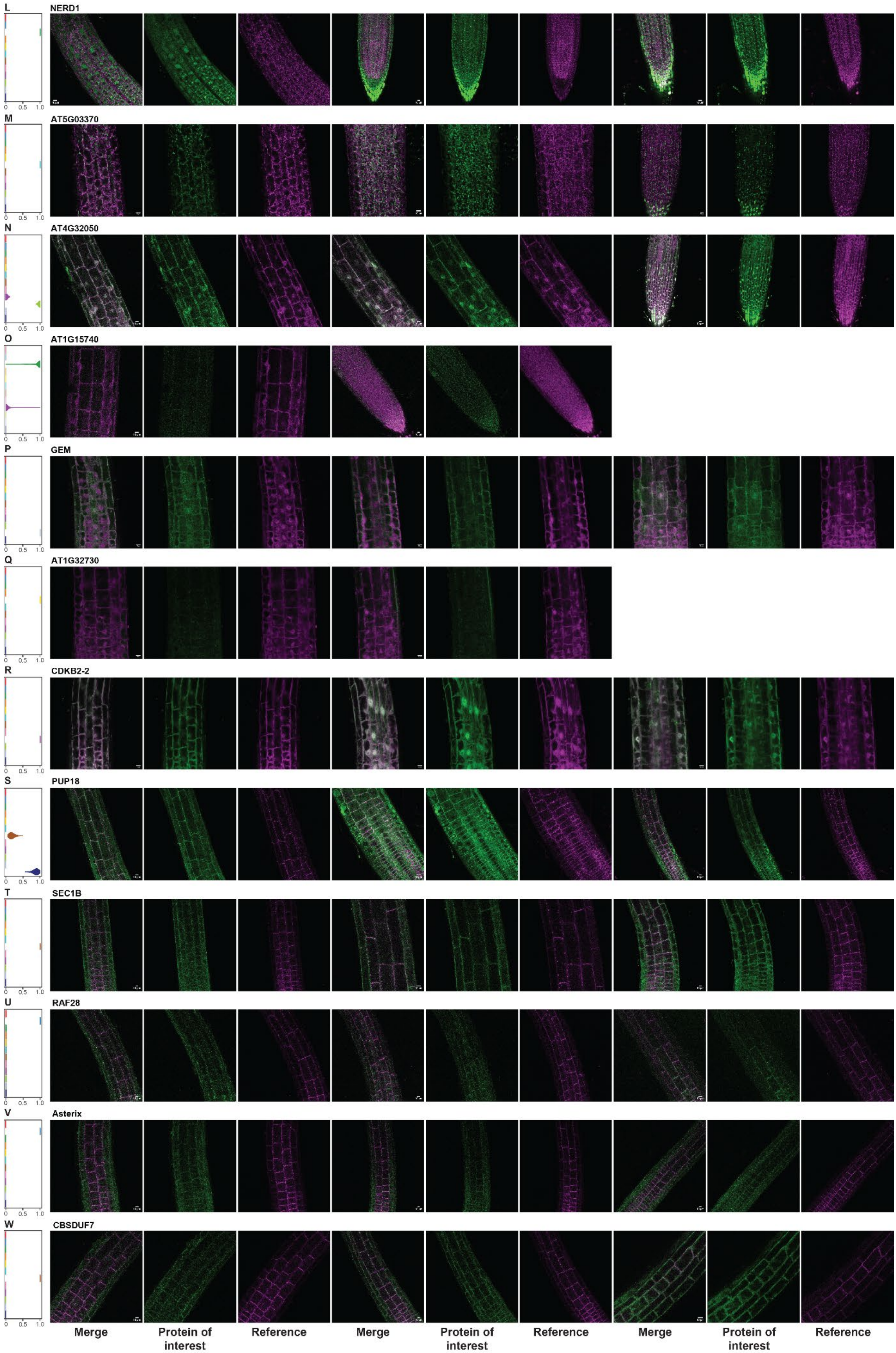
L
NERD1
M
AT5G03370
N
AT4G32050
O
AT1G15740
P
GEM
Q
AT1G32730
R
CDKB2-2
S
PUP18
T
SEC1B
U
RAF28
V
Asterix
W
CBSDUF7
0
0.5
1.0
Merge
Protein of interest
Reference
Merge
Protein of interest
Reference
Merge
Protein of interest
Reference

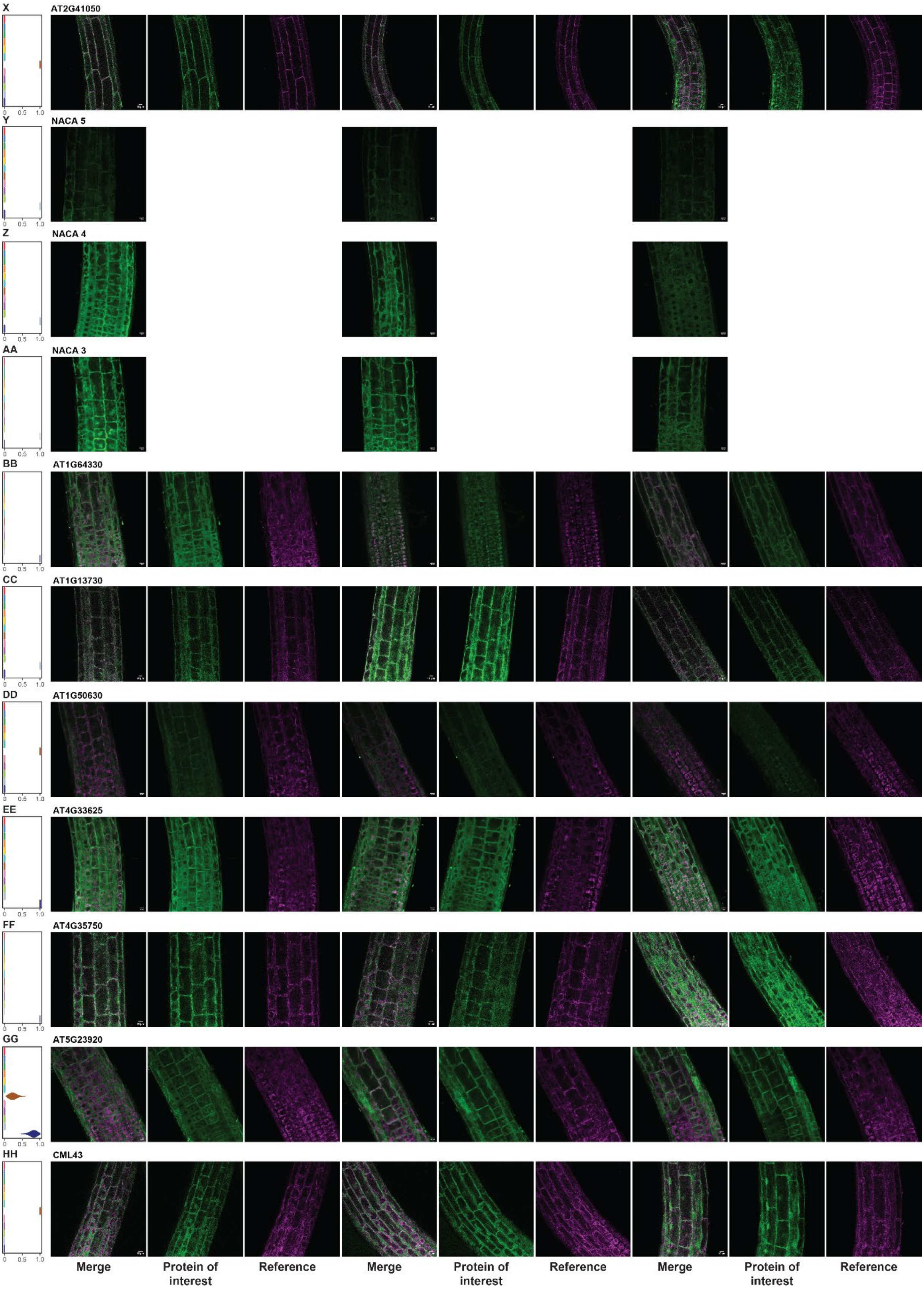


**Figure S4: Annotations to subcellular localizations and confocal imaging for a selection of 35 proteins; related to Figure 3.** (A-HH) Annotation of subcellular localization and confocal imaging to confirm localizations *in vivo*. Details on the lines can be found in Table S1.

**Table S1: overview of the validation of the Arabidopsis root map by confocal microscopy; related to Figure 3.** In the table predictions, background lines and the number of matches to the annotated localization are described.

| Araport | Uniprot | Gene name | TAGM-MCMC prediction | Background line | Publication background line | Match |
|---|---|---|---|---|---|---|
| **AT5G61760** | Q9FLT2 | IPK2B | Cytosol | WAVE-1R free RFP | Geldner et al. 2009[44] | 11 of 11 |
| **AT5G04120** | F4KI56 | AT5G04120 | Cytosol | WAVE-1R free RFP | Geldner et al. 2009[44] | ? |
| **AT2G43980** | O80568 | IPTK4 | Cytosol | WAVE-1R free RFP | Geldner et al. 2009[44] | 10 of 10 |
| **AT2G47970** | O82264 | NPL42 | Cytosol | WAVE-1R free RFP | Geldner et al. 2009[44] | 8 of 8 |
| **AT5G07960** | Q9SD88 | ASTERIX | ER | WAVE-138 PM | Geldner et al. 2009[44] | ? |
| **AT4G31170** | Q9M085 | RAF28 | ER | WAVE-138R PM | Geldner et al. 2009[44] | ? |
| **AT5G42570** | Q93XZ7 | AT5G42570 | ER | WAVE-6R ER | Geldner et al. 2009[44] | 5 of 5 |
| **AT2G37540** | O80924 | AT2G37540 | ER | WAVE-6R ER | Geldner et al. 2009[44] | 7 of 7 |
| **AT1G71780** | F4IA30 | AT1G71780 | ER | WAVE-6R ER | Geldner et al. 2009[44] | 6 of 6 |
| **AT5G56980** | F4K956 | PAMPA70 | ER | WAVE-6R ER | Geldner et al. 2009[44] | 5 of 5 |
| **AT4G00670** | Q6IDB4 | AT4G00670 | ER | WAVE-6R ER | Geldner et al. 2009[44] | 0 of 6 |
| **AT2G15240** | Q9SKL4 | AT2G15240 | Golgi | WAVE-22R Golgi | Geldner et al. 2009[44] | 0 of 0 |
| **AT1G50120** | A0A1P8ASX9 | LOT | Golgi | WAVE-22R Golgi | Geldner et al. 2009[44] | 10 of 10 |
| **AT3G51050** | F4J381 | NERD1 | Golgi | WAVE-22R Golgi | Geldner et al. 2009[44] | 0 of 10 |
| **AT2G47650** | Q8S8T4 | UXS4 | Golgi | WAVE-22R Golgi | Geldner et al. 2009[44] | 7 of 7 |
| **AT1G15740** | Q8H1Q4 | AT1G15740 | Golgi (95) Plastid Ribosomal (5) | WAVE-1R free RFP | Geldner et al. 2009[44] | ? |
| **AT1G31730** | Q9LPJ1 | AT1G32730 | Ribosome | WAVE-1R free RFP | Geldner et al. 2009[44] | ? |
| **AT5G03370** | Q9LZF2 | AT5G03370 | Peroxisome | Mit-GFP | Nelson et al. 2007[45] | 10 of 10 |
| **AT1G20930** | Q8LG64 | CDKB2-2 | Plastid ribosomal | WAVE-1R free RFP | Geldner et al. 2009[44] | 5 of 5 |
| **AT1G47330** | Q8RY60 | CBSDUF7 | PM | WAVE-138 PM | Geldner et al. 2009[44] | 11 of 11 |
| **AT2G41050** | Q8RXY4 | AT2G41050 | PM | WAVE-138R PM | Geldner et al. 2009[44] | 8 of 10 |

| | | | | | | |
|---|---|---|---|---|---|---|
| **AT4G12120** | Q9SZ77 | SEC1B | PM | WAVE-138R PM | Geldner et al. 2009[44] | 11 of 11 |
| **AT5G44460** | Q9FI19 | CML43 | PM | WAVE-9 Vacuole | Geldner et al. 2009[44] | 0 of 10 |
| **AT1G50630** | Q9LPT3 | AT1G50630 | PM | WAVE-9 Vacuole | Geldner et al. 2009[44] | 13 of 21 |
| **AT4G32050** | Q5E911 | AT4G32050 | Proteasome (99) | WAVE-1R free RFP | Geldner et al. 2009[44] | 8 of 8 |
| **AT5G13850** | Q6ICZ8 | NACA3 | Ribosome | Col-0 | N/A | 5 of 5 |
| **AT4G10480** | Q9SZY1 | NACA4 | Ribosome | Col-0 | N/A | 5 of 5 |
| **AT1G33040** | Q8LGC6 | NACA5 | Ribosome | Col-0 | N/A | 7 of 7 |
| **AT1G13730** | Q9LMX6 | AT1G13730 | Nucleus | WAVE-13 TGN | Geldner et al. 2009[44] | ? |
| **AT2G22475** | Q8S8F8 | GEM | Ribosome | WAVE-1R free RFP | Geldner et al. 2009[44] | 5 of 5 |
| **AT4G33625** | Q8GXK1 | AT4G33625 | Vacuole | WAVE-9 Vacuole | Geldner et al. 2009[44] | 12 of 12 |
| **AT4G35750** | O81806 | AT2G37540 | ER | WAVE-9 Vacuole | Geldner et al. 2009[44] | 9 of 9 |
| **AT1G64330** | Q9C7V7 | AT1G64330 | Vacuole | WAVE-9 Vacuole | Geldner et al. 2009[44] | 17 of 17 |
| **AT5G23920** | Q9FF88 | AT5G23920 | Vacuole (77) PM (23) | WAVE-9 Vacuole | Geldner et al. 2009[44] | 7 of 12 |
| **AT1G57990** | Q9C508 | PUP18 | Vacuole (83) PM (16) | WAVE-138R PM | Geldner et al. 2009[44] | 8 of 8 |

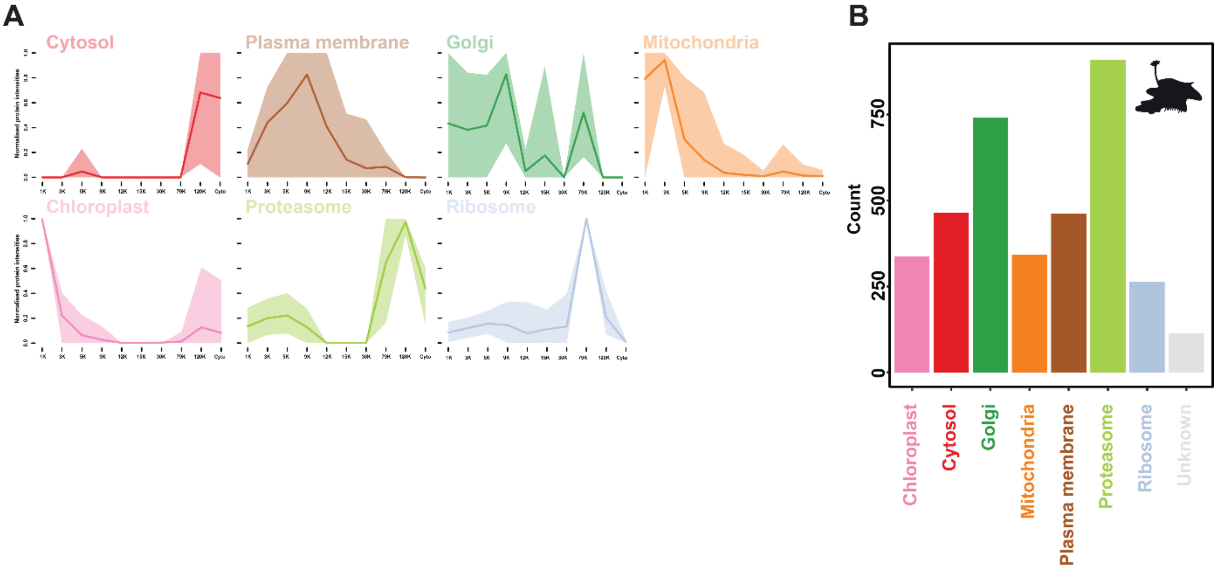


**Figure S5: Obtained fractionation profiles of marker proteins and distribution of proteins over subcellular localizations for Marchantia; related to Figure 4.** (A) Fractionation patterns for the marker proteins for every subcellular localization in the Marchantia dataset. (B) Distribution of the number of annotated proteins for every subcellular localization in the Marchantia dataset

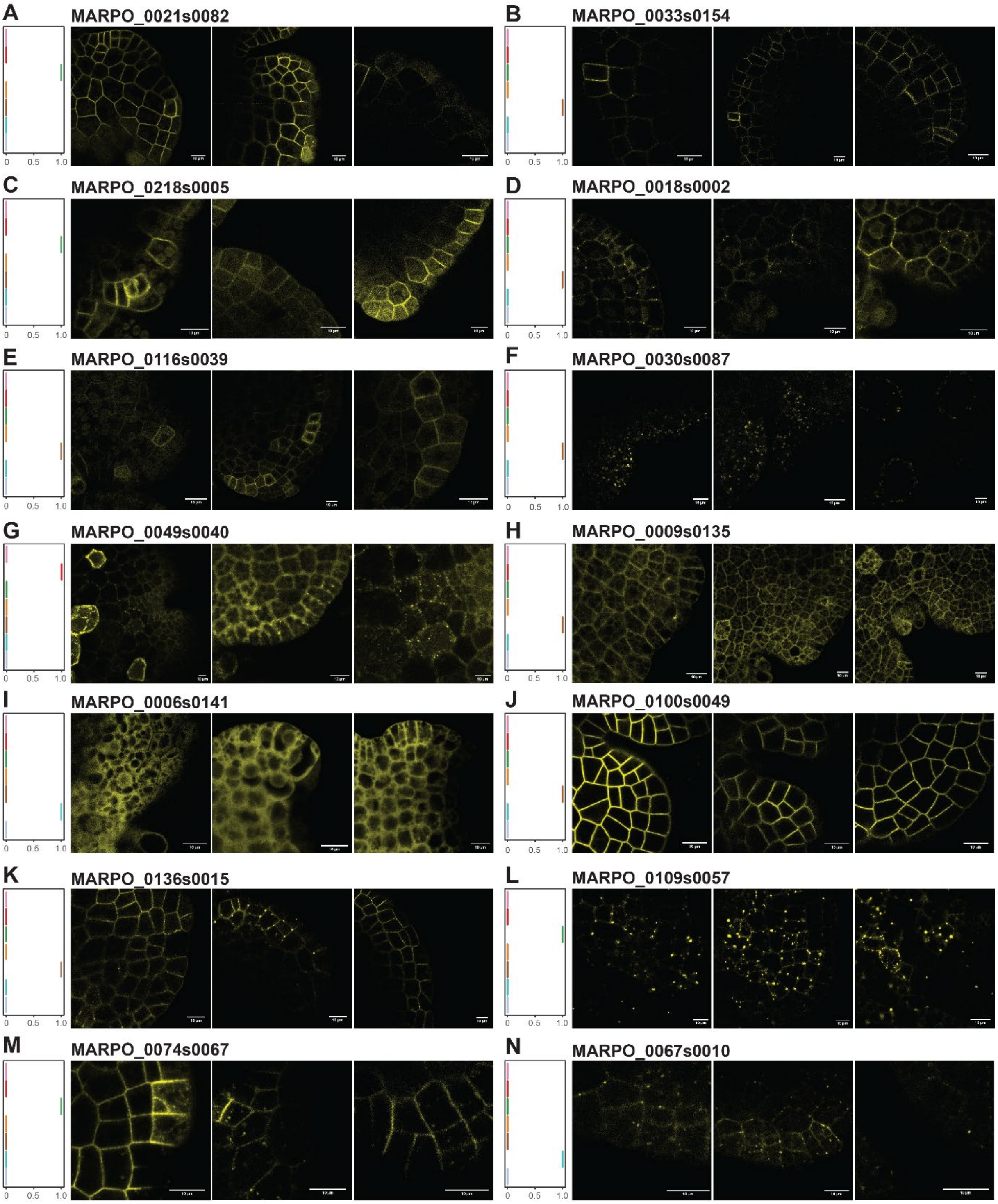


**Figure S6: Annotations to subcellular localizations and confocal imaging for a selection of 35 proteins; related to Figure 4.** (A-N) Annotation of subcellular localization and confocal imaging to confirm localizations *in vivo*. Details on the lines can be found in Table S2.

**Table S2: overview of the validation of the Arabidopsis root map by confocal microscopy; related to Figure 4.** In the table predictions and the number of matches to the annotated localization are described.

| MARPO_# | Uniprot | TAGM-MCMC prediction | Match |
|---|---|---|---|
| **MARPO_0074s0067** | A0A1B4Z1I7 | Golgi | Maybe |
| **MARPO_0067s0010** | A0A2R6WQ21 | Peroxisome | Yes |
| **MARPO_0018s0002** | A0A2R6VWR4 | Plasma membrane | Yes |
| **MARPO_0100s0049** | A0A2R6WEQ5 | Plasma membrane | Yes |
| **MARPO_0021s0082** | A0A2R6XDP6 | Golgi | No |
| **MARPO_0049s0040** | A0A2R6WY14 | Cytosol | Yes |
| **MARPO_0033s0154** | A0A2R6X6P5 | Plasma membrane | Yes |
| **MARPO_0136s0015** | A0A1B4Z1K0 | Plasma membrane | Yes |
| **MARPO_0009s0135** | A0A2R6XLQ9 | Plasma membrane | Yes |
| **MARPO_0116s0039** | A0A2R6WB29 | Plasma membrane | Yes |
| **MARPO_0109s0057** | A0A2R6WCS5 | Golgi | Yes |
| **MARPO_0218s0005** | A0A2R6VZU3 | Golgi | No |
| **MARPO_0006s0141** | A0A2R6XPS0 | Peroxisome | No |
| **MARPO_0030s0087** | A0A2R6X8B9 | Plasma membrane | No |

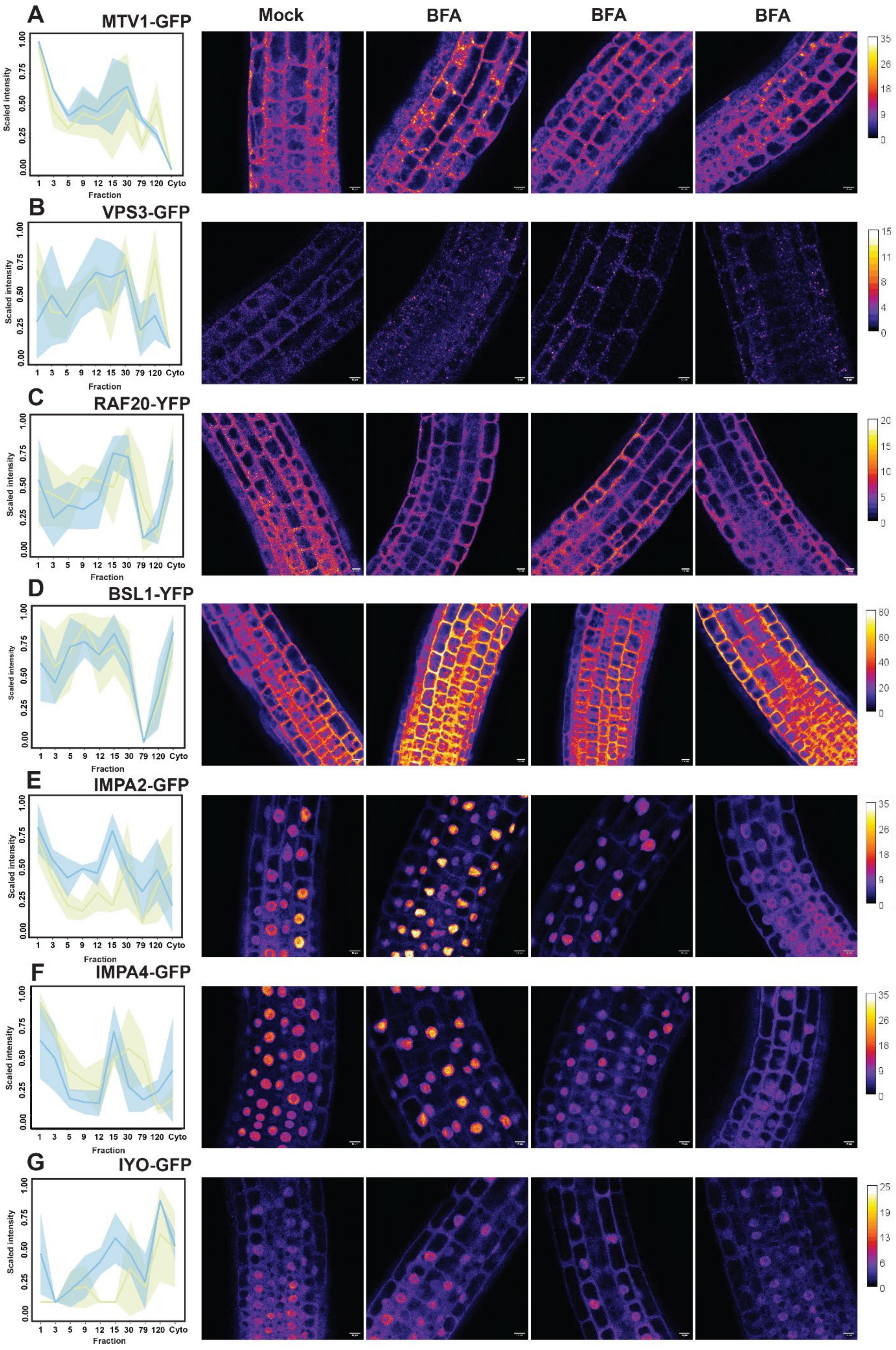

A
MTV1-GFP
B
VPS3-GFP
C
RAF20-YFP
D
BSL1-YFP
E
IMPA2-GFP
F
IMPA4-GFP
G
IYO-GFP
Mock
BFA
BFA
BFA
Scaled intensity
Fraction
1 3 5 9 12 15 30 79 120 Cyto
0.00 0.25 0.50 0.75 1.00

**Figure S7: Confocal imaging of 1 hour 10 µM BFA treatment on Arabidopsis roots; related to Figure 5.** (A-G) Fractionation patterns of the protein in mock (blue) and BFA-treated (green) conditions and confocal imaging. Details are described in Table S3.

**Table S3: Overview of the validation of the BFA treatment by confocal microscopy following a 1 hour 10 µM treatment; related to Figure 5.** In the table, the original publication, the observed effect of the treatment and the number of affected plants is described.

| Line | Publication | Araport | UniProt | Effect | nr of affected plants |
|---|---|---|---|---|---|
| **BSL1** | Guo et al., 2021[114] | AT4G03080 | Q8L7U5 | None | 0 of 8 |
| **IMPA2** | Bhattacharjee et al., 2008[70] | AT4G16143 | F4JL11 | Altered nucleus cytoplasm ratio | 9 of 9 |
| **IMPA4** | Bhattacharjee et al., 2008[70] | AT1G09270 | O80480 | Altered nucleus cytoplasm ratio | 8 of 12 |
| **IYO** | Munoz et al., 2017[72] | AT4G38440 | Q8GYU3 | None | 0 of 10 |
| **MTV1** | Sauer et al., 2013[69] | AT3G16270 | Q9C5H4 | Potential BFA bodies | 7 of 9 |
| **RAF20** | Kuhn et al., 2024[53] | AT1G79570 | Q9SAJ2 | Loss of puncta | 14 of 17 |
| **VPS3** | Takemoto et al., 2018[113] | AT1G22860 | F4I312 | Potential BFA bodies | 9 of 10 |
